# Myhre syndrome variants form long-lived SMAD heterotrimers through a rewired polar interface at the variant site

**DOI:** 10.64898/2026.11.20.768049

**Authors:** Carles Torner, Radoslaw Pluta, Miriam Condeminas, Rameez J. Khan, Marina R. de Régil, Lidia Ruiz, Joan Pous, Israel Ramos, María Caballero, Anne-Sophie Humm, Stephan Niebling, Maria García-Alai, Jose A. Márquez, Aurora Martinez, Simon Fromm, Pau Martin-Malpartida, Maria J. Macias

**Affiliations:** Institute for Research in Biomedicine (IRB Barcelona), The Barcelona Institute of Science and Technology (BIST). Baldiri Reixac, 10, Barcelona 08028, Spain; European Molecular Biology Laboratory, Grenoble, EMBL, 71, Avenue des Martyrs, CS 90181 38042 Grenoble Cedex 9, France; European Molecular Biology Laboratory, Notkestrasse 85, Hamburg, Germany; Department of Biomedicine, and Neuro-SysMed, Department of Neurology, Haukeland University Hospital, 5020 Bergen, Norway; European Molecular Biology Laboratory, Meyerhofstraße 1 69117 Heidelberg, Germany; ICREA, Passeig Lluís Companys 23, 08010-Barcelona, Spain

## Abstract

TGF-β signaling through SMAD transcription factors governs development and tissue homeostasis. Most disease-associated SMAD4 mutations cause loss of function, but five germline variants at only two positions in the MH2 domain, Arg496 and Ile500, instead cause Myhre syndrome, a rare condition inducing pathway dysregulation. We show that the two most common Myhre variants form more stable heterotrimers with SMAD3 than wild-type SMAD4, with affinities increasing two-to threefold and complex half-lives lengthening about fivefold. Cryo-EM reconstructions and twelve crystal structures show that a few interfacial hydrogen bonds shorten to 2.5 Å, reorganizing the binding interface. Our models also identified the SMAD4 C-terminus bound oriented resembling a phosphorylated R-SMAD tail, and two non-equivalent SMAD3 N-termini observed in the absence of cofactors. To the extent that these structural features reflect the cellular environment, they provide a mechanistic basis for the altered signaling dynamics observed in Myhre syndrome and outline concrete avenues for pharmacologically targeting these stabilized complexes.

## Introduction

Germline substitutions at two recurrent positions in SMAD4 cause Myhre syndrome (MYHRS, OMIM #139210) <u>(Myhre et al. 1981)</u>. This rare congenital disorder is characterized by progressive fibrosis of the skin, lungs, heart, and gastrointestinal tract, alongside connective and skeletal tissue anomalies, cardiovascular stenosis, speech and motor delays, and immune dysfunction (reviewed in <u>(Pindon et al. 2025)</u>). Initially, these clinical features were documented primarily in childhood; however, a recent cohort study of 24 affected adults—17 of whom were diagnosed after age 18—extends the phenotypic spectrum into adulthood and identifies scoliosis as a previously unrecognized manifestation <u>(Vanbelleghem et al. 2024; Cappuccio et al. 2022; Le Goff et al. 2011; Pindon et al. 2025; Lin et al. 2020; Caputo et al. 2012; Yang et al. 2022)</u>. As with many rare diseases, clinical management remains limited to symptomatic relief.

The great majority of disease-associated SMAD4 variants act in the opposite direction and they reduce or abolish signaling and contribute to cancer onset and progression <u>(Massagué and Sheppard 2023; Macias et al. 2015)</u>. The Myhre-associated variants instead resist the regulatory steps that normally terminate signaling. In patient-derived nasal epithelial cells carrying I500V, R496C or I500T, SMAD4 levels and upstream R-SMAD phosphorylation are unchanged while transcriptional activity is increased, an effect attributed to a stronger protein–protein interaction within the SMAD complex <u>(Lindsay et al. 2025)</u>. However, this hypothesis has not been tested on the complex itself.

SMAD4 is the common mediator of transforming growth factor-beta (TGF-β) family signaling, a pathway that integrates inputs from a large superfamily of cytokines and membrane receptors and engages in extensive cross-talk with other signaling pathways. As a result, the network governs embryonic development, tissue homeostasis, immune maturation and neurovascular regulation <u>(Luo 2017; Massagué 2012; Wang et al. 2014)</u>. Ligand binding activates receptor kinases that phosphorylate receptor-regulated SMADs (R-SMADs), which then assemble with SMAD4 into heterotrimeric complexes that translocate to the nucleus and regulate gene expression (simplified in **Figure 1a**) <u>(Kawabata et al. 1998; Jayaraman and Massague 2000; Saka et al. 2007)</u>. SMAD4 and the R-SMADs share a modular architecture with two domains connected by a dynamic linker that allows SMAD proteins to adopt closed and open conformations and modulates the SMAD interaction with activators and repressors <u>(Gomes et al. 2021)</u>. SMADs have an N-terminal MH1 domain that binds DNA <u>(Jatzlau et al. 2025;Martin-Malpartida et al. 2017; Ruiz et al. 2021)</u>, a C-terminal MH2 domain that mediates trimerization and cofactor binding, and a flexible linker carrying docking sites for kinases and ubiquitin ligases <u>(Shi and Massagué 2003; Alarcón et al. 2009; Gao et al. 2009; Aragón et al. 2011; Macias et al. 2015)</u>.

**Figure 1.**
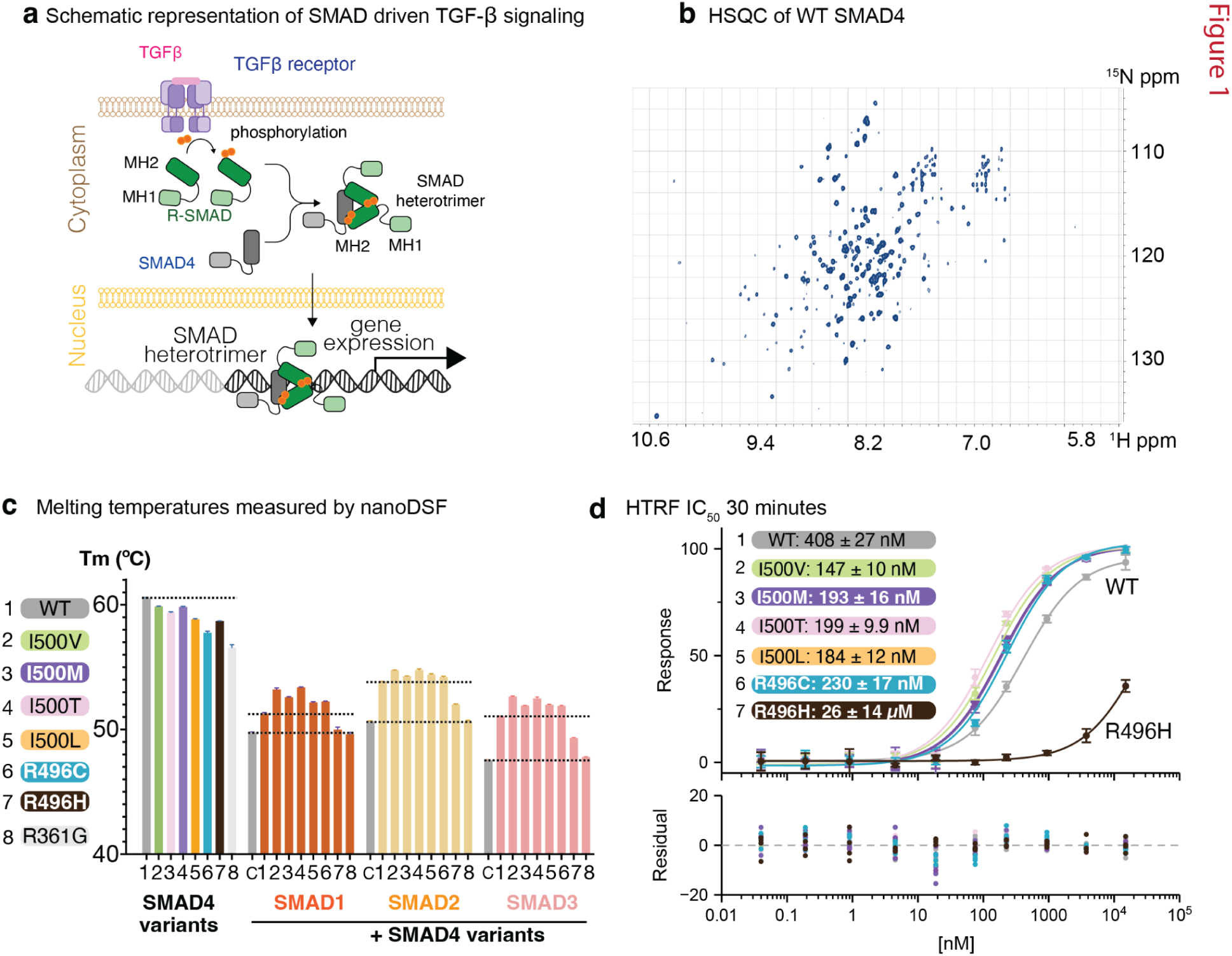
Myhre syndrome variants preserve the SMAD4 MH2 fold and enhance its interaction with R-SMADs. a, Schematic of SMAD-mediated TGF-β signaling. Ligand binding activates TGF-β receptors, which phosphorylate the C-terminal motif of receptor-regulated SMADs (R-SMADs). Phosphorylated R-SMADs form heterotrimers with SMAD4 through their MH2 domains; the complexes then enter the nucleus and bind DNA through their MH1 domains to regulate gene expression. MH1 and MH2 domains are indicated b, ¹H–¹⁵N HSQC spectrum of the WT SMAD4 MH2 domain (starting at amino acid 314) 900 MHz, 298K. Spectra of the R496C and I500V variants are shown in **Supplementary Figure 1b**. c, Melting temperatures (T_m_) measured by nanoDSF. Left, SMAD4 MH2 WT and variants 1–8 alone. Right, phosphomimetic SMAD1, SMAD2 and SMAD3 MH2 domains alone, ‘C’, or mixed with each SMAD4 variant (1–8). Dotted lines mark the Tm of WT SMAD4, of each R-SMAD alone and of each R-SMAD with WT SMAD4. Bars show mean ± s.d.; n = 4 technical replicates. d, HTRF competition assay measuring inhibition of the SMAD3–SMAD4 complex signal by increasing concentrations of SMAD4 MH2 WT and variants. Measurements were made after 30 min incubation. Lines are fits to a three-parameter logistic model. The inset gives IC50 values with S.E. Data are mean ± s.d.; n = 4 technical replicates.

The lifetime of the assembled complex is itself regulated. In the nucleus, TRIM33 and SMURF2 monoubiquitinate SMAD4 at Lys519 to impede its association with phosphorylated R-SMADs and to drive its nuclear export <u>(Luo et al. 2026; Dupont et al. 2009)</u>, and SMURF2 monoubiquitinates SMAD3 at Lys333 and Lys378 <u>(Tang et al. 2011)</u>, whereas acetylation of SMAD3 Lys378 by p300/CBP has the opposite effect of stimulating SMAD-driven transcription <u>(Inoue et al. 2007)</u>. Lowering TRIM33, for example by DCAF6-mediated degradation, reduces SMAD4 monoubiquitination and increases SMAD2/3–SMAD4 association and target-gene output in cells <u>(Luo et al. 2026)</u>. In parallel, inhibitory SMADs compete with SMAD4 for binding to activated R-SMADs <u>(Hata et al. 1998; Itoh et al. 2008; Miyazawa and Miyazono 2017)</u>. Each of these steps depends on the assembly state of the complex, yet how a pathogenic variant might evade these regulatory mechanisms has not been examined structurally.

MYHRS variants are confined to two positions within the SMAD4 MH2 domain and comprise five recurrent substitutions—R496C and the I500V/M/T/L cluster—with R496C and I500V accounting for nearly 87% of reported cases <u>(Lin et al. 2024; Caputo et al. 2012; Alankarage et al. 2022; Wood et al. 2024; Caputo et al. 2014; Le Goff et al. 2011)</u>. Despite introducing markedly different physicochemical perturbations such as side-chain volume reduction (I500V and I500T), methyl rearrangement (I500L), insertion of a flexible thioether (I500M), or charge removal (R496C), these substitutions all result in the same clinical syndrome, suggesting they converge on a shared molecular mechanism. Structurally, however, the two positions play distinct roles: Arg496 resides on helix H4 and directly contacts the R-SMAD in the wild-type heterotrimer, whereas Ile500, located on strand β10, is buried within SMAD4 and makes no direct R-SMAD contact, instead packing directly against Arg496. Intriguingly, other residues along this same interface including Asp493, Arg497, and Glu526, located only a few angstroms away are also mutated in human disease, yet those substitutions cause loss of function. Thus, a single interaction surface appears capable of tuning TGF-β signaling in opposite directions, though the structural determinants separating these two outcomes remain unknown.

To test this hypothesis, we combined isothermal titration calorimetry (ITC), binding kinetics, differential scanning calorimetry (nano-DSF), and single-molecule mass measurements with high-resolution structural analyses. Specifically, we determined single-particle cryo-electron microscopy (cryo-EM) reconstructions of nanobody-bound SMAD4–SMAD3 heterotrimers alongside crystal structures of the wild-type MH2 domain and all five MYHRS variants, in which a crystallographic three-fold axis reproduces the R-SMAD-binding interface. Together, these results show that the five substitutions indeed converge on a shared molecular mechanism. Every variant stabilizes the complex with SMAD3, the two variants analyzed kinetically form longer-lived complexes, and the overall dimensions of the interface are not systematically altered; instead, a small number of polar contacts are re-organized relative to the WT, shifting binding from an entropy-driven to an enthalpy-dominated regime for the I500 substitutions. The cryo-EM reconstructions additionally allow us to trace terminal segments and loops left disordered in previous crystal structures, including the lysins through which the complex is normally switched off, and resolve coexisting conformations that reveal how the two R-SMAD subunits differ in a given complex.

## Results

### 1. Myhre substitutions at two SMAD4 positions converge on longer-lived SMAD3 heterotrimers

#### 1.1 Myhre variants retain the MH2 fold and DNA binding but marginally destabilize the isolated domain

We first establish the biophysical consequences of the Myhre variants (**Figures 1 and 2**), then define their structural basis at one interface (**Figures 3 and 4**), and finally describe additional features of SMAD3–SMAD4 complexes revealed by the structures (**Figure 5**). Because the Myhre variants lie in the SMAD4 MH2 domain, we first asked whether they alter its tertiary structure (constructs, tags and theoretical masses are shown in **S_Figure 1a**). Two-dimensional NMR spectra of the two most prevalent clinical variants, I500V and R496C, show chemical-shift dispersion closely similar to the wild-type (WT) domain, indicating that both are correctly folded; in all three spectra the resonances differ in line shape and intensity, as expected for a domain containing both well-ordered elements and dynamic loops (**Figure 1b, S_Figure1b**). We then tested whether, in the context of the full-length protein, the variants might promote inhibitory contacts with the MH1 domain and so restrict access to DNA. However, electrophoretic mobility shift assays with full-length WT, I500V and R496C proteins showed no detectable difference in DNA binding (**S_Figure 1c**).

**Figure 2.**
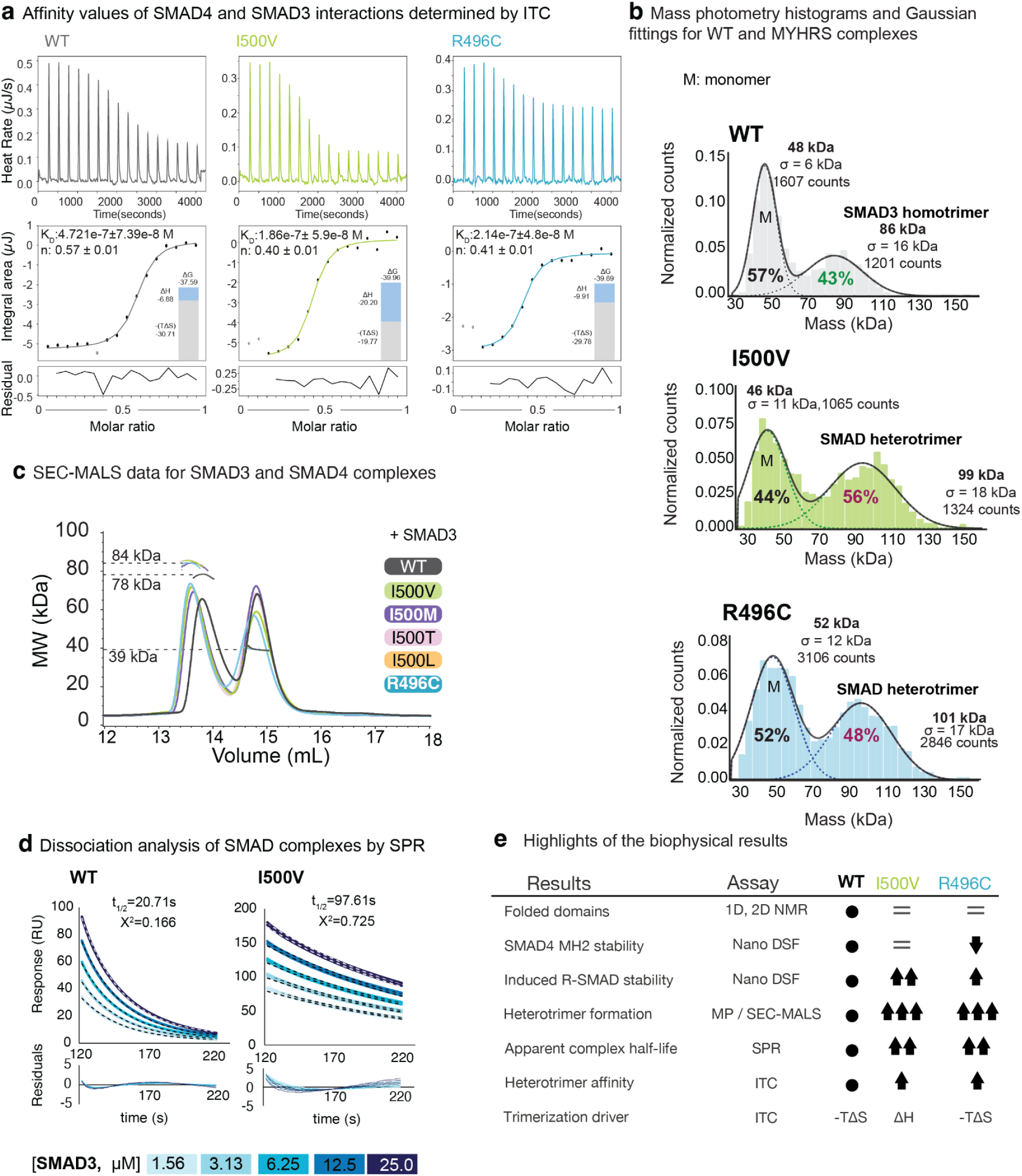
SMAD4 variants I500V and R496C enhance heterotrimer formation with SMAD3 and slow its dissociation. (a) Isothermal titration calorimetry (ITC) of SMAD3 binding to WT SMAD4 (grey), I500V (green), and R496C (blue) measured at 37 °C. Top, raw thermograms. Middle, integrated heats fitted to a one-site binding model, with the fitted K_D_ and stoichiometry (n) shown in each panel. Insets show the thermodynamic signature (ΔG, ΔH, and −TΔS). Bottom, fit residuals. Grey points were excluded from the fit. Thermodynamic values are typically reported per mole in kJ/mol. (b) Mass photometry histograms of SMAD3 mixed with WT SMAD4, I500V, or R496C. Solid lines show the overall Gaussian fits and dotted lines the individual components. Peak masses, standard deviations (σ), counts, and relative populations are indicated. M, monomer. For WT, the species at ∼86 kDa corresponds to the SMAD3 homotrimer, whereas for I500V (99 kDa) and R496C (101 kDa) the larger species corresponds to the SMAD heterotrimer. (c) Size-exclusion chromatography with multi-angle light scattering (SEC-MALS) of SMAD3 combined with WT SMAD4 or the variants I500V, I500M, I500T, I500L, and R496C (colors as indicated). Molecular weight (MW) across the elution peaks is shown for the WT (78 kDa) and variant (84 kDa) complexes. The 39 kDa dashed line marks the monomeric species. (d) Surface plasmon resonance (SPR) dissociation analysis of SMAD3 complexes with WT SMAD4 (left) and I500V (right). The R496C variant is shown as Supplementary Figure 2d. SMAD3 was injected at 1.56–25.0 µM (colors as indicated). Black dashed lines show the dissociation model fits, with the apparent half-life (t₁/₂) and χ² indicated. Residuals are shown below each sensorgram. Dissociation of the I500V complex is markedly slower than that of WT (t₁/₂ = 97.61 s versus 20.71 s). (e) Summary of the biophysical results for I500V and R496C relative to WT, accross assays. Arrows indicate the direction and relative magnitude of the change, and "=" indicates no change. The thermodynamic driver of trimerization is shown as entropy-driven (−TΔS) for WT and R496C and enthalpy-driven (ΔH) for I500V.

**Figure 3.**
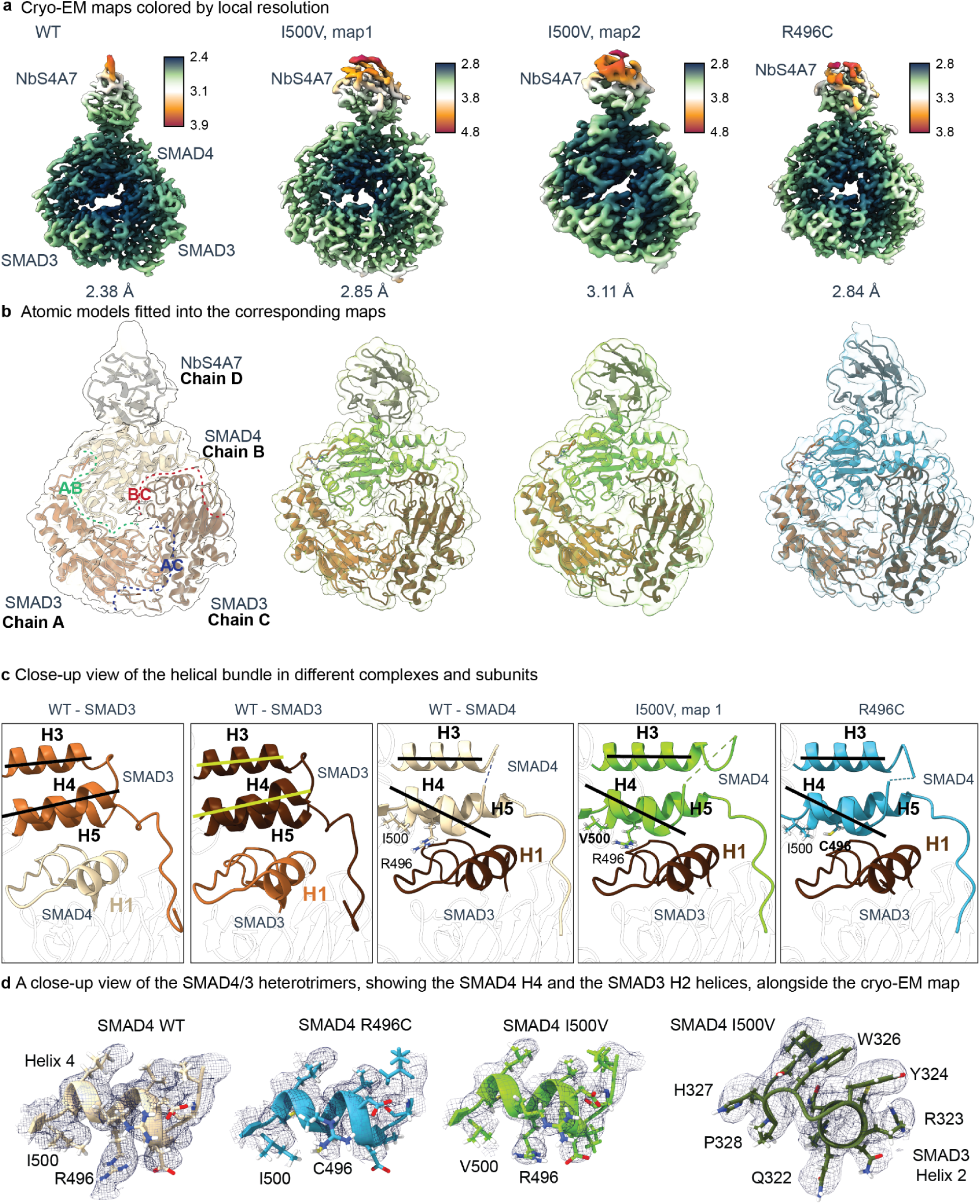
Cryo-EM structures of NbS4A7-bound SMAD4–SMAD3 WT and variant heterotrimers. (a) Cryo-EM maps of NbS4A7–SMAD4 MH2–2x(SMAD3 MH2) complexes containing SMAD4 WT (2.38 Å), I500V map 1 (2.85 Å), I500V map 2 (3.11 Å) and R496C (2.84 Å), colored by local resolution (Å; scale bar beside each map). Global resolutions are from gold-standard FSC = 0.143. (b) Atomic models fitted into the corresponding maps (transparent surfaces). Chains A and C, SMAD3; chain B, SMAD4; chain D, NbS4A7. Dashed outlines on the WT model mark the B–A, A–C and B–C subunit interfaces. These models are depicted with hydrogens for illustration purposes. (c) Close-up views of the three-helix bundle (H3–H5) packing against helix H1 of the neighbouring subunit. Shown are the two SMAD3 subunits and the SMAD4 subunit of the WT complex, the SMAD4 subunit of I500V class 1 and the SMAD4 subunit of R496C. Lines mark the axes of helices H3 and H4 and show the different tilt of H4 (32°) and H3 (14°) in SMAD4 compared with SMAD3. Residues 496 and 500 are shown as sticks. Cryo-EM processing workflows are shown in **Schematics 1 and 2**. Domain topologies and sequence comparison are shown in **Supplementary Figure 3**. (d) Close-up views of the SMAD4 helix H4 and SMAD3 helix H2, shown with the corresponding cryo-EM density (grey mesh). From left to right, SMAD4 WT, R496C and I500V, each showing the side chains at positions 496 and 500. Right, the same region of the I500V map showing the SMAD3 helix H2 residues Q322, R323, Y324, W326, H327 and P328 that pack against it.

**Figure 4.**
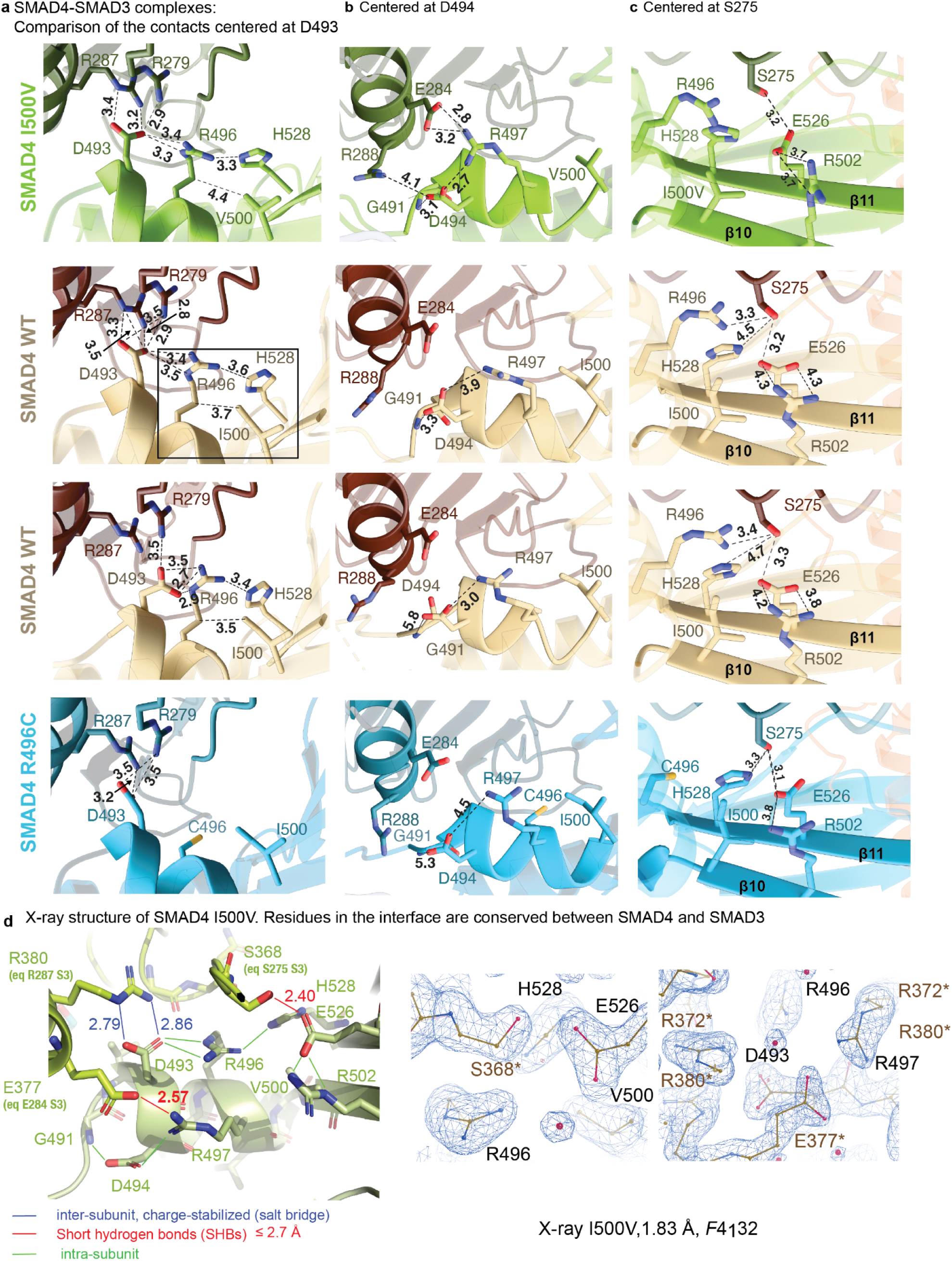
Myhre variants rewire polar contacts at the SMAD4–SMAD3 interface. (a–c) Inter-subunit contacts centred on SMAD4 D493 (a), SMAD4 D494 (b) and SMAD3 S275 (a) in the cryo-EM structures of the I500V (top), WT (midle) and R496C (bottom) complexes. SMAD3 is shown in darker shades. Dashed lines indicate polar contacts with their distances. The box highlights the R496–I500 in WT contact (also dashed line, but it is not a HB). (b) Crystal structure of SMAD4 MH2 I500V (1.83 Å; space group F4₁32), in which the homotrimer is generated by the crystallographic three-fold axis. Interface residues are conserved between SMAD4 and SMAD3; equivalent SMAD3 residues are given in parentheses. Line colours: blue, inter-subunit salt bridges; red, short hydrogen bonds (≤2.7 Å); green, intra-subunit contacts. Right, 2Fo–Fc electron density (contoured at 1σ) for the I500V structure. Asterisks mark residues from the neighboring subunit.

**Figure 5.**
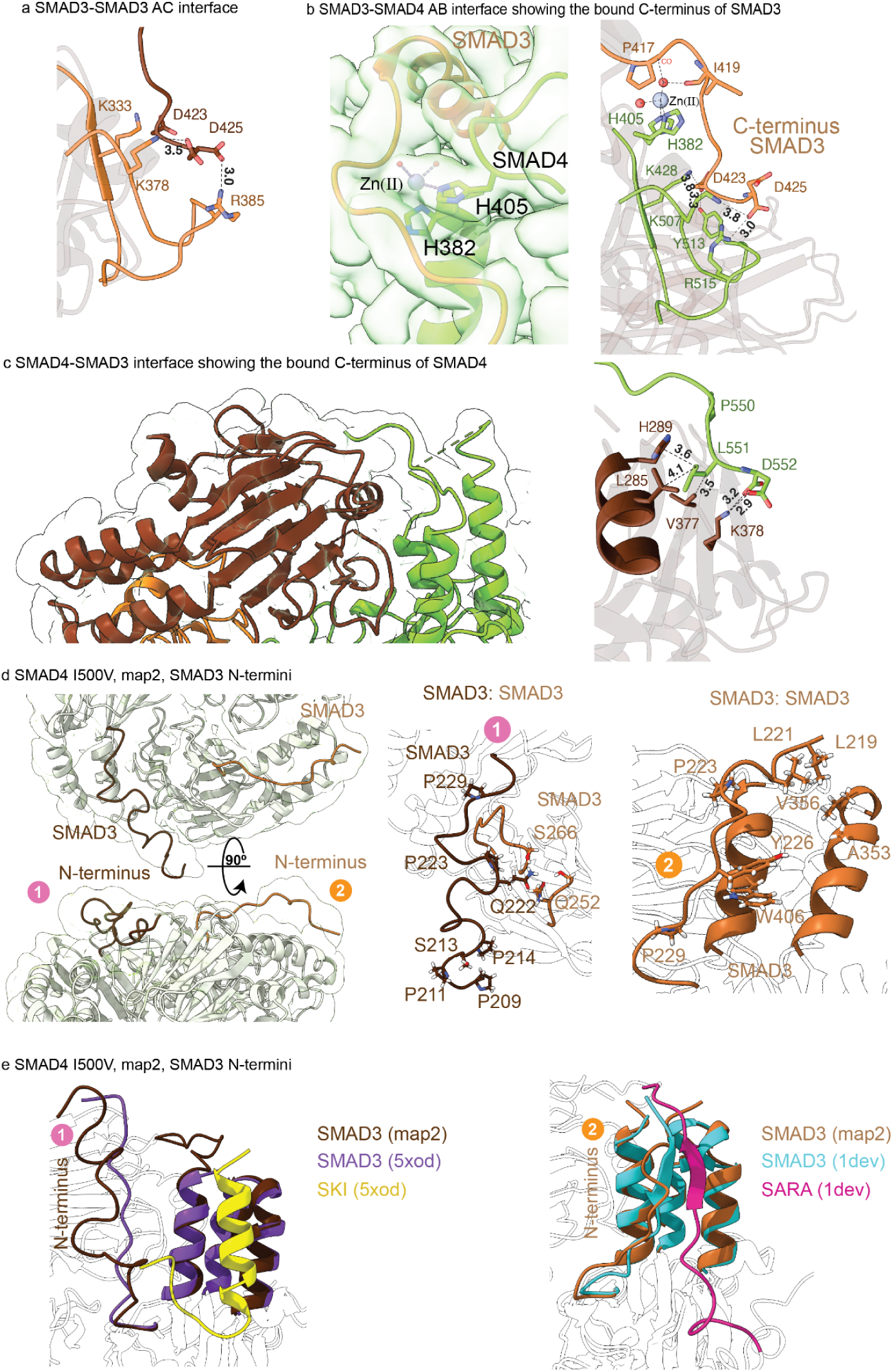
Terminal segments stabilize the SMAD4–SMAD3 and SMAD3–SMAD3 interfaces. (a) The SMAD3– SMAD3 A–C interface in the I500V map 1 structure. The C-terminal D423 and D425 of one SMAD3 subunit engage K333, K378, and R385 of the neighbouring subunit. (b) The SMAD4–SMAD3 A–B interface, showing the bound phosphomimetic C-terminus of SMAD3. Left, map and model showing a putative Zn(II) ion coordinated by SMAD4 H382 and H405. Right, SMAD3 D423 and D425 contacting SMAD4 K428, K507 and R515.(c) The SMAD4–SMAD3 interface in the I500V map 1 structure, showing the C-terminus of SMAD4 (green) bound to SMAD3. Right, close-up of SMAD4 P550, L551 and D552 packing against SMAD3 R385 and R386 near helix H1. (d) Ordered SMAD3 N-terminal segments (residues 219–229) in the I500V class 2 map. Left, two views related by a 90° rotation, with the two N-termini labeled 1 and 2. Centre and right, close-ups of each N-terminus and its contacts with the neighbouring SMAD3 subunit and its own SMAD3, respectively. (e) Superposition of N-terminus 1 with SMAD3 bound to SKI (PDB 5XOD) and of N-terminus 2 with SMAD3 bound to SARA (PDB 1DEV).

#### 1.2 All five variants stabilize R-SMADs and outcompete wild-type SMAD4

Since DNA binding was retained, we turned to the assembly of heterotrimeric complexes with R-SMADs. We monitored the thermal stability (T_m_) of the phosphomimetic SMAD1, SMAD2 and SMAD3 MH2 domains in the presence of increasing concentrations of each SMAD4 variant by nanoDSF (differential scanning fluorimetry with light scatering). This approach is feasible because the baseline T_m_ values of isolated R-SMADs are approximately 10°C lower than that of WT SMAD4 <u>(Martin-Malpartida et al. 2022; Martin-Malpartida et al. 2024)</u>. The isolated variant domains were only marginally less stable than WT (T_m_ lower by 0.5–2.5 °C, against 3.8 °C for the loss-of-function control R361G), leaving sufficient thermal separation from the R-SMADs for a robust titration (**Figure 1c)**. To optimize the titration conditions, we performed a pilot experiment with WT SMAD4 and SMAD3, observing a concentration-dependent increase in SMAD3 T_m_ upon SMAD4 binding. Applying 2:1 R-SMAD:SMAD4 stoichiometry ratio to SMAD1, SMAD2 and SMAD3, every R-SMAD was stabilized more by each Myhre variant than by WT SMAD4. For SMAD3, WT SMAD4 raised the T_m_ from 47.5 to 51.1 °C, whereas the five variants reached 51.9 to 52.7 °C; R361G left the SMAD3 T_m_ near that of SMAD3 alone and R496H raised it by only 1.8 °C (**Figure 1c**, T_m_ values). To test this effect independently we performed competitive FRET-based homogeneous time-resolved fluorescence (HTRF) assays using SMAD3 and SMAD4 variants. Starting with preformed, high-FRET WT-SMAD4/SMAD3 heterotrimers, we titrated increasing concentrations of unlabeled SMAD4 variants to displace the WT protein. In agreement with the DSF observations, the addition of MYHRS variants resulted in a concentration-dependent decrease in the FRET signal in all cases, with IC_50_ values of 147 to 230 nM for variants and 26 µM for R496H compared with 408 nM for WT (**Figure 1d**), thereby confirming that SMAD3 forms more stable complexes with the variants than with the WT.

#### 1.3 Higher affinity accounts for part of the effect

We first asked how much affinity alone contributes. ITC of each SMAD4 variant against phosphomimetic SMAD3 gave dissociation constants from 75 nM (I500M) to 214 nM (R496C), compared with 472 nM for WT SMAD4 - a two-to sixfold gain across the five Myhre variants (**Figure 2a, S_Figure 2a**). Fitted stoichiometries ranged from 0.40 to 0.57, bracketing the 0.5 expected for a 1:2 SMAD4:SMAD3 assembly. Substitutions at position I500 also shifted the thermodynamic signature as binding became more enthalpically favorable by 9 to 15 kJ/mol relative to WT, suggesting that new polar contacts might form at the interface, with a compensating entropic cost. A two-to sixfold change in affinity is, however, too small on its own to explain a qualitative change in complex abundance, which led us to ask how the complexes behave kinetically.

#### 1.4 I500V and R496C complexes dissociate 4.7-fold more slowly

Equilibrium affinity cannot distinguish faster assembly from slower dissociation, so we measured dissociation directly by surface plasmon resonance, with WT, I500V and R496C SMAD4 amine-coupled to separate channels of one sensor chip and SMAD3 injected as analyte. Fitting the dissociation phase to a single-exponential model gave complex half-lives of 97.6 s for I500V and 98.2 s for R496C against 20.7 s for WT (**Figure 2d, S_Figure 2d**), a 4.7-fold extension of residence time and a larger effect than the corresponding change in affinity. Because only the dissociation phase was analyzed, the association rate was not determined; the longer residence time is nonetheless sufficient to account for the increased stability of the variant complexes.

#### 1.5 Variant heterotrimers persist after dilution at nM and µM concentrations

Because the measured affinities span the nanomolar to micromolar range, we assessed the abundance and oligomeric state of the complexes at both ends of this range by mass photometry and by size-exclusion chromatography coupled to multi-angle light scattering (SEC-MALS). This question has both functional implications and practical consequences for the preparation of heterotrimers for structural studies.

At nanomolar concentrations, mass photometry of nanobody-labeled mixtures detected no heterotrimer with WT SUMO-SMAD4, whereas the heterotrimer accounted for 56% (I500V) and 48% (R496C) of the particles counted for the variants (**Figure 2b, S_Figure 2b**). At micromolar concentrations, SEC-MALS carried out without the nanobody and at a 2:1 SMAD4:SMAD3 ratio, chosen so that SMAD3 is driven into heterotrimer rather than into its own homotrimer, resolved the heterotrimer as a prominent, monodisperse peak for all five variants and as only a minor species for WT (**Figure 2c, S_Figure 2c**). Across three orders of magnitude in concentration, therefore, the variants shift the equilibrium from free SMAD4 and SMAD3 homotrimers toward the assembled heterotrimer.

The enhanced stability reflects higher binding affinity combined with a prolonged residence time, increasing the population of heterotrimeric signaling complexes at both nanomolar and micromolar concentrations (summarized in **Figure 2e**).

### 2. The variants rewire polar contacts at a single interface

#### 2.1 Structures at side-chain resolution define the wild-type and variant interfaces

The biophysical characterization of the variants indicates that a localized variant introduces changes large enough to extend complex lifetime 4.7-fold, yet small enough to leave the overall fold, the NMR fingerprint and the thermal stability of the isolated domain essentially unchanged. We therefore needed structures of the assembled complexes at a resolution sufficient to compare side-chain geometry, together with crystal structures of the isolated domains at a resolution sufficient to measure hydrogen-bond distances and specific details.

We determined cryo-EM structures of SMAD3 heterotrimers with wild-type SMAD4 and with the I500V and R496C variants. These complexes are demanding targets for single-particle analysis. as they are relatively small (approx. 87 kDa), disc-shaped, and with an apparent threefold pseudo-symmetry that complicates particle classification. We addressed this using a SMAD4-specific nanobody (NbS4A7) whose binding site does not affect the interaction of SMAD4 with the variants or with R-SMADs <u>(Condeminas et al. 2026)</u>, but breaks the apparent pseudo-symmetry, increases the particle size to ∼99 kDa, and facilitates particle classification. Homogeneous refinement yielded maps at 2.38 Å (WT), 2.85 Å (I500V, map 1) and 2.84 Å (R496C) (**Figure 3a,b**; **Table 1**). In each model chain B is SMAD4, chains A and C are SMAD3 and chain D is the nanobody, so that the two SMAD4-SMAD3 interfaces are B-A and B-C and the SMAD3-SMAD3 interface is A-C. These maps also cover regions not captured in previous crystal structures, such as the SMAD4 C-terminal extension and the loops carrying the lysins through which the complex is regulated.

**Table 1.** Cryo-EM data collection, processing and refinement statistics.

| **SMAD3 complexes with SMAD4** | **WT** | **I500V-MAP1** | **I500V-MAP2** | **R496C** |
|---|---|---|---|---|
| PDB entry | 50OY | 50OZ | 50PA | 50PB |
| EMDB entry | EMD-100814 | EMD-100815 | EMD-100816 | EMD-100817 |
| **Data collection and processing** |  |  |  |  |
| Magnification | 215,000 | 130,000 | 130,000 | 165,000 |
| Electron exposure (e<sup>-</sup>/Å<sup>2</sup>) | 60 | 60 | 60 | 60 |
| Defocus range (µm) | 0.6 to 2.2 | 0.5 to 1.5 | 0.5 to 1.5 | 0.5 to 2.5 |
| Pixel size (Å; map voxel size) | 0.5726 | 0.934 | 0.934 | 0.731 |
| Symmetry imposed | C1 | C1 | C1 | C1 |
| Initial particle images (no.) | 3,046,741 | 7,353,612 | 7,353,612 | 3,905,268 |
| Final particle images (no.) | 478,661 | 958,007 | 32,648 | 287,322 |
| Map resolution range (Å) | 2.4-4.1 | 2.67-3.93 | 2.67-3.93 | 2.64-3.80 |
| **Refinement** |  |  |  |  |
| Model resolution (Å) | 2.38 | 2.85 | 3.10 | 2.84 |
| Non-hydrogen atoms | 5,598 | 5,728 | 5,764 | 5,663 |
| ● Protein | 5,598 | 5,711 | 5,763 | 5,660 |
| ● Ligands / ions | 0 | 1 | 1 | 1 |
| ● Water | 0 | 2 | 0 | 2 |
| Protein residues | 708 | 727 | 730 | 716 |
| Mean B factor (Å<sup>2</sup>) |  |  |  |  |
| ● Protein | 51 | 48 | 88 | 46 |
| ● Ligands / ions | N.A. | 80 | 156 | 87 |
| ● Water | N.A. | 52 | N.A. | 54 |
| R.m.s. deviations: bond lengths (Å) | 0.002 | 0.002 | 0.002 | 0.001 |
| R.m.s. deviations: bond angles (°) | 0.43 | 0.43 | 0.47 | 0.41 |
| **Validation** |  |  |  |  |
| MolProbity Score | 1.16 | 1.06 | 1.39 | 1.18 |
| Clashscore | 3.00 | 1.51 | 2.91 | 1.98 |
| Rotamer outliers (%) | 0.00 | 0.00 | 0.00 | 0.00 |
| Ramachandran favoured (%) | 97.70 | 97.07 | 95.56 | 96.60 |
| Ramachandran allowed (%) | 2.30 | 2.93 | 4.44 | 3.40 |
| Ramachandran outliers (%) | 0.00 | 0.00 | 0.00 | 0.00 |
| Model-to-map CC (mask) | 0.81 | 0.86 | 0.77 | 0.80 |

Although the final consensus reconstructions resolved the structured cores at high resolution including the variant sites and all main loops, certain regions remained highly heterogeneous, and conformational averaging left the density corresponding to several flexible areas at the N- and C-termini to become weak or absent. To characterize this heterogeneity and recover structural signals obscured during homogeneous refinement, in the I500V–SMAD3 complex we returned to the particle datasets and performed three-dimensional variability analysis (3DVA). Along the resulting variability trajectories, density corresponding to the C-terminal tail of SMAD4-MH2 and the N-terminal tails of SMAD3-MH2 progressively appeared and disappeared, consistent with these regions sampling different conformations and degrees of structural order, shown in **Supplementary Video 1**. Guided by these observations, we performed focused three-dimensional classification of the relevant regions, followed by non-uniform refinement of the selected particle classes. This strategy separated a distinct conformational state that we analyzed and allowed us to obtain a second I500V map (map 2) at 3.11Å. Thus, each I500V-SMAD3 map shows distinct interpretable densities for the terminal segments, allowing us to trace additional residues and to identify previously unresolved inter-subunit contacts. The processing workflows are illustrated in **Schematics 1, 2.**

We also determined twelve crystal structures of the isolated SMAD4 MH2 domain (**Table 2, Supplementary Table 1**), in addition to one previously determined of the WT (PDB 9HWE). Eleven structures belong to space group F4₁32 and were refined at resolution of 1.73–2.13 Å, comprising three independent crystals of I500V, four of R496C, two of I500M, and one each of I500T and I500L. Each structure contained a single molecule in the asymmetric unit and assembled into a homotrimer about the crystallographic three-fold. As a result, packing is constant throughout the series and differences can be assigned to the substitution rather than to the lattice. A twelfth structure, of the WT in space group P6₃ at 2.06 Å, determines the same contacts in a different lattice and under different crystallization conditions. The crystallographic homotrimer reproduces the heterotrimer contact surface closely, as the same thirteen partner positions engage the Myhre patch, and the residues forming the contacts analyzed here, Ser368′, Glu377′ and Arg380′, are identical to their SMAD3 counterparts Ser275, Glu284 and Arg287. However, the contact surface is not equivalent as SMAD4 Leu381and SMAD3 Arg288 are different, so interactions involving that position cannot be assessed in the crystals. Moreover, crystal packing restrains the inter-subunit motion that the reconstructions reveal, so the homotrimer reports interface geometry at a fixed point rather than the range sampled in solution. Replicate crystals separate reproducible geometry from refinement noise: Glu526–Ser368′, the contact on which our comparison rests, varies by no more than 0.12 Å between crystals of the same variant, a fifth of its difference from the WT. Ordered molecules from the condition never lie within 3.70 Å of residue 496 or 500, and in each of the seven cases where an additive lies within 4 Å of a contact used in our comparisons, a crystal lacking that additive reproduces the same geometry.

**Table 2.** X-ray data collection, processing and refinement statistics.

| **Structure** | **SMAD4 I500V** | **SMAD4 I500T** | **SMAD4 I500L** | **SMAD4 I500M** | **SMAD4 R496C** | **SMAD4 WT** |
|---|---|---|---|---|---|---|
| **PDB ID** | 35RN | 35RO | 35RP | 35RQ | 35RR | 35RS |
| **Data collection** |  |  |  |  |  |  |
| Beamline | ALBA XALOC | ALBA XALOC | ALBA XALOC | ALBA XALOC | ALBA XALOC | ALBA XALOC |
| Wavelength (Å) | 0.97926 | 0.97926 | 0.97926 | 0.97926 | 0.97926 | 0.97926 |
| Space group | F 41 3 2 | F 41 3 2 | F 41 3 2 | F 41 3 2 | F 41 3 2 | P 63 |
| Unit cell | 195.79 195.79 195.79<br>90 90 90 | 195.71 195.71 195.71<br>90 90 90 | 196.22 196.22 196.22<br>90 90 90 | 196.66 196.66 196.66<br>90 90 90 | 195.98 195.98 195.98<br>90 90 90 | 97.33 97.33 115.86<br>90 90 120 |
| Resolution shells (Å) | overall, inner, outer | overall, inner, outer | overall, inner, outer | overall, inner, outer | overall, inner, outer | overall, inner, outer |
|  | 56.52 – 1.83, 56.52 – 5.27, 1.88 - 1.83 | 69.19 – 1.80, 69.19 – 5.18, 1.85 - 1.80 | 27.48 – 1.73, 27.48 – 4.69, 1.76 - 1.73 | 59.30 – 1.78, 59.30 – 5.11, 1.84 - 1.78 | 69.29 – 1.84, 69.29 – 5.31, 1.89 - 1.84 | 84.29 – 2.06, 84.29 - 6.00, 2.21 - 2.06 |
| *R*<sub>merge</sub> | 0.083, 0.027, 1.955 | 0.051, 0.019, 1.804 | 0.118, 0.039, 8.556 | 0.066, 0.026, 2.049 | 0.050, 0.023, 1.402 | 0.061, 0.028, 1.367 |
| *R*<sub>pim</sub> | 0.027, 0.009, 0.596 | 0.017, 0.007, 0.584 | 0.014, 0.005, 0.961 | 0.019, 0.008, 0.571 | 0.020, 0.010, 0.528 | 0.029, 0.014, 0.530 |
| *I*/s(*I*) | 16.6, 52.0, 1.4 | 22.9, 73.9, 1.4 | 22.7, 129.7, 0.3 | 21.1, 65.1, 1.4 | 18.8, 57.2, 1.5 | 16.3, 45.9, 1.7 |
| Wilson B-factor | 35.62 | 37.33 | 37.48 | 35.64 | 38.58 | 50.63 |
| Completeness (%)* |  |  |  |  |  |  |
| - spherical | 95.5, 99.6, 60.8 | 95.5, 99.7, 56.0 | 100.0, 100.0, 100.0 | 95.5, 99.0, 54.4 | 94.5, 99.0, 57.4 | 81.7, 97.5, 22.1 |
| - ellipsoidal | 95.5, 99.6, 60.7 | 95.5, 99.7, 56.0 | Not applicable | 95.5, 99.0, 54.4 | 94.4, 99.0, 57.3 | 94.1, 97.5, 67.5 |
| Multiplicity | 10.6, 9.2, 11.5 | 9.7, 8.4, 10.4 | 77.0, 69.4, 79.5 | 13.3, 11.1, 13.7 | 6.9, 6.1, 8.0 | 5.2, 4.9, 7.6 |
| *CC*<sub>1/2</sub> | 0.999, 1.000, 0.516 | 1.000, 1.000, 0.527 | 1.000, 1.000, 0.281 | 1.000, 1.000, 0.574 | 0.999, 0.999, 0.565 | 0.999, 0.999, 0.536 |
| **Refinement** |  |  |  |  |  |  |
| Resolution (Å)<sup>#</sup> | 56.52 - 1.83 (1.9 - 1.83) | 48.93 - 1.80 (1.86 - 1.80) | 27.48 - 1.73 (1.78 - 1.73) | 59.29 - 1.78 (1.85 - 1.78) | 69.29 - 1.84 (1.90 - 1.84) | 47.74 - 2.06 (2.13 - 2.06) |
| Reflections used in refinement<sup>#</sup> | 27509 (1807) | 29042 (1775) | 34150 (2718) | 30224 (1778) | 27023 (1679) | 31229 (440) |
| Reflections used for R-free<sup>#</sup> | 1387 (90) | 1420 (91) | 1727 (138) | 1464 (90) | 1353 (78) | 1542 (31) |
| *R*<sub>work</sub> / *R*<sub>free</sub> (%) | 19.9/22.3 | 19.2/21.8 | 19.9/21.0 | 20.2/21.5 | 19.8/22.2 | 18.3/20.0 |
| Protein residues | 201 | 201 | 191 | 195 | 192 | 349 |
| B factors | 50.49 | 53.00 | 57.50 | 49.64 | 53.66 | 62.87 |
| RMS(bonds) | 0.008 | 0.005 | 0.010 | 0.007 | 0.008 | 0.003 |
| RMS(angles) | 0.97 | 0.80 | 1.12 | 0.99 | 0.93 | 0.64 |
| Ramachandran |  |  |  |  |  |  |
| Favored (%) | 97.46 | 96.95 | 95.72 | 94.76 | 96.81 | 97.63 |
| Outliers (%) | 0.51 | 0.51 | 0.53 | 0.52 | 0.53 | 0.00 |
| Rotamer Outlies (%) | 0.00 | 0.60 | 1.22 | 0.00 | 0.00 | 0.00 |
| Clashscore | 3.12 | 3.13 | 3.25 | 6.14 | 1.31 | 1.41 |

\* Anisotropy correction by STARANISO/autoPROC with the default setting used for the determination of the resolution cutoff (Tickle et al. 2018; Vonrhein et al. 2011); ellipsoidal completeness is given when anisotropy correction applied. <sup>#</sup> Statistics for the highest-resolution shell are shown in parentheses.

#### 2.2 The SMAD4 helical bundle is the most dynamic element of the heterotrimer

The MH2 fold is a compact alpha/beta domain of roughly 200 residues that forms the C-terminal half of R-SMAD and SMAD4 proteins. The topology of SMAD3 and SMAD4 MH2 domains in the cryo-EM structure of the WT SMAD4/SMAD3 complex are indicated in **S_Figure 3a** and the sequence comparison of both domains in **S_Figure 3b**. As we and others previously observed in X-ray structures (9HWE, 1YGS) <u>(Condeminas et al. 2026; Macias et al. 2025; Shi et al. 1997)</u>, the domain core consists of two curved beta sheets packed face-to-face in a sandwich arrangement with a larger sheet of six strands (2,1,11,10,7,8) and a smaller one of five strands (4, 3, 5, 6 and 9). The β-sandwich has a complex topology, with strand connectivity between the two sheets interleaved rather than sequential. The sandwich is flanked by five α-helices (H1–H5) and one conserved 3₁₀ helix in the L3 loop (**S_Figure 3a**). In all subunits, helices H1 and H2 pack against the outer β-sheet face, while the C-terminal three-helix bundle (H3–H5) caps the opposite side to form the heterotrimer interfaces. While this bundle is less dynamic and fully traced in SMAD3, it exhibits marked conformational variability in SMAD4 (**Supplementary Video 1**). We attribute this variability to a confluence of two features involving H3 and H4 helices in the bundle. First, in isolated SMAD4 crystal structures <u>(Condeminas et al. 2026; Chacko et al. 2004)</u>, H3 can have different lengths depending on the conditions, extending up to 12 residues further due to crystal packing. In contrast, our heterotrimer cryo-EM maps reveal a compact helical bundle of comparable size across subunits, with H3 slightly longer in the I500V variant and shifted with respect to the orientation in SMAD3 by 14°(map 1). Second, in SMAD4, H4 is rotated by 32° relative to its SMAD3 counterpart as reflected in the superimposition of the different helical bundles of the SMAD proteins (**Figure 3c**).

This altered orientation, combined with steric hindrance from the perpendicular SMAD3 H1 helix in the heterotrimeric complex, and the trajectory of the SMAD4 C-terminus when bound to SMAD3, prevents H4 from extending prior to residue I490 without creating clashes with H3 and H5. Consequently, a ∼20-amino-acid segment in SMAD4 spanning part of H3 and the H3–H4 loop remains partially unresolved due to dynamic conformational exchange. 3DVA captures this heterogeneity, which is consistent with a role in heterotrimerization as the SMAD4 C-terminal tail engages SMAD3 (**Supplementary Video 1**). Close-up views of the cryo-EM maps with tracing the side chains for WT and for the I500V and R496C variants are included as **Figure 3d**.

#### 2.3 Myhre substitutions rewire the polar network at the β10 and H4-H1 contact

The MH2 heterotrimer is built from three equivalent interfaces, at each of which one subunit donates the face formed by helices H4 and H5 and its C-terminal segment, and the neighboring subunit receives it with loop L2, helix H1, the β3–β4 edge and the basic pocket of the L3 loop. In the cryo-EM heterotrimers, the SMAD4-SMAD3 interface carrying the substitutions is formed between chains B and C, and is defined by strands β10 and helix H4 of SMAD4 against helix H1 of SMAD3 (dotted lines in **Figure 3b**). In the wild-type complex, SMAD4 Arg496 and Ile500 sit in close proximity, held in place by His528, contacts from SMAD3 Ser275 to Glu526, and an intra-subunit Asp494–Arg497 interaction that stabilizes helix H4 (**Figure 4a,b middle and S_Figure 4d**). Adjacent to Arg496, Asp493 samples two conformations and acts as a central anchoring point, coordinating either three arginine side chains (SMAD3 Arg279 and Arg287 across the interface together with SMAD4 Arg496) or, in the alternative conformer, just Arg279 and Arg496. In crystal structures of the isolated domain, an ordered water molecule forms a HB with the Arg496 guanidinium in eight of the nine structures that retain Arg496.

Replacing the bulky Ile500 side chain with valine creates a cavity and makes the hydrophobic packing against Arg496 less efficient (**Figure 4a, top**). The interface responds not by contracting as a whole but by re-forming its polar contacts locally. For instance, in the I500V-SMAD3 complex, the SMAD4 backbone shifts along strand β10 and helix H4 toward SMAD3, helix H3 lengthens by one turn relative to the wild type, and side chains that were not engaged in contacts in the wild type, come into hydrogen-bonding or electrostatic contact range. SMAD4 Arg497 reaches SMAD3 Glu284 and SMAD4 Asp494 does so with SMAD3 Arg288, neither of which is in contact in the WT map, while SMAD4 Asp493 (only one rotamer) maintains its contact with SMAD3 Arg287.

In contrast to the conservative substitutions at position 500, R496C replaces a long basic side chain with a compact uncharged one, reducing contacts with adjacent residues (**Figure 4a bottom**). Whereas Asp493 contacts SMAD3 Arg287 (3.2 Å) in the cryo-EM map, the Asp493–Arg380′ interaction is lost in the R496C crystal structures: although three of the four crystals contain an ordered additive 2.85–3.31 Å from this pair, the fourth—free of additives within 9 Å—shows the same separation (5.28 Å), demonstrating that the rearrangement arises from the substitution rather than crystallization conditions. R496C does, however, retain the short Glu526–Ser275 contact in the heterotrimer (**Figure 4c, bottom**) as well as its equivalent Glu526–Ser368′ hydrogen bond in the crystals (2.54 Å). Thus, the R496C heterotrimer exhibits no new contacts at the Myhre patch and approximately half as many close contacts between SMAD4 β10/H4 and SMAD3 H1 compared to the WT or I500V complexes. This difference is matched by the ITC experiments. The I500 substitutions shift binding to an enthalpy-dominated regime (ΔH −15.9 to −21.8 kJ/mol against −6.9 for the WT), consistent with consolidating several polar contacts at once, whereas R496C remains entropy-dominated like the WT (−9.9 kJ/mol) (**S_Figure 2a**), consistent with an exchange that gains as much as it loses. Thus, substitutions at position 500 and R496C reach a similar endpoint, tighter binding, by two structural mechanisms with distinct thermodynamic signatures.

We also asked whether these changes alter the overall dimensions of the assembly. All values use the residue set modeled in every reconstruction (SMAD3 residues 227–425 in both subunits, SMAD4 residues 318–543 and 111 nanobody residues), with the nanobody interface as internal reference. On this basis the four complexes are almost not distinguishable (**S_Table 2**). The variants therefore alter specific interfacial contacts without a major alteration of the overall proximity of the subunits.

Because the changes in the cryo-EM maps are too small to quantify differences in distances reliably, we measured the equivalent contacts in the crystal structures of the SMAD4 alone, where the resolution supports it. Among the comparable contacts (**S_Figure 4b**), there is one short contact that behaves identically in all five variants (between the carboxylate of Glu526 and the hydroxyl of Ser368′). This contact measures 2.4–2.6 Å (+/- 0.06 Å) across the eleven variant crystals, against 2.8-2.9 Å in the WT crystals (**S_Figure 4c-h**). As a reference, the equivalent contact in the heterotrimer, SMAD4 Glu526 to SMAD3 Ser275, is present in the cryo-EM models of the variants (**Figure 4c**).

#### 2.4 Variants that remove these specific contacts fail to stabilize SMAD3 upon binding

To test whether these contacts hold the SMAD complexes together, we evaluated a panel of disease- and cancer-associated SMAD4 variants from cBioPortal—rather than standard alanine scanning—with SMAD3 as the binding partner, using nanoDSF to monitor complex stability. This panel includes SMAD4 substitutions located directly at the SMAD4–SMAD3 interface, in its immediate vicinity, or at distant structural sites. This panel includes SMAD4 substitutions located directly at the SMAD4–SMAD3 Myhre syndrome interface, in its immediate vicinity, or at distant structural sites. All isolated SMAD4 domains in the panel were properly folded and thermally stable. Alone, SMAD3 displays a melting temperature T_m_ of 47.5 °C, which increases upon SMAD4 binding to 51.1 °C with the wild-type (WT) protein and to 51.9–52.7 °C with the Myhre syndrome variants (**S_Figure 5a**). Substitutions that leave SMAD3 essentially unstabilized are inferred to abolish efficient binding: for instance, SMAD3 reaches a T_m_ of 47.8 °C with R361G and 48.5 °C with E526K, both within 1.0 °C of isolated SMAD3. Conversely, substitutions introducing an alternative hydrogen-bonding side chain retain partial stabilization, raising the SMAD3 T_m_ to 50.5 °C with R497H and 49.3 °C with R496H. The sharpest contrast occurs at a single position: mutating Arg496 to cysteine raises the SMAD3 T_m_ by 0.8 °C relative to WT, whereas mutating it to histidine lowers the T_m_ by 1.8 °C, a 2.6 °C spread dictated by side-chain chemistry rather than sequence position. In contrast, A406T, which makes no contact with SMAD3 in the structures, shows only a slight decrease in stabilization relative to WT SMAD4 (T_m_ = 50.0 °C). Thus, other disease-associated substitutions located at the Myhre syndrome interface—namely R496H, R497H, and E526K, where specific contacts are rewired by Myhre mutations—consistently reduce the thermal stabilization of SMAD3.

### 3. The structures reveal new features of SMAD3–SMAD4 complexes

#### 3.1 A Zn(II) site formed by two SMAD4 histidines distinguishes the two SMAD3 C-termini

In the cryo-EM heterotrimers of WT and I500V, the three C-terminal segments bind cyclically, with each subunit donating its C-terminus to the basic pocket of the adjacent subunit and receiving the C-terminus of the preceding one. The two SMAD3 subunits are therefore non-equivalent: the C-terminus of SMAD3 chain A is read by SMAD4, whereas the C-terminus of chain C is read by the other SMAD3 subunit, and SMAD4 in turn donates its own C-terminus to chain C—although the receiving pocket occupies the same structural site in both proteins. Each pocket comprises two sub-sites that separately accommodate the two phosphorylatable positions of an R-SMAD tail: position 423 engages the tyrosine hydroxyl and lysine of one sub-site, while position 425 engages the arginine and lysine of the other (**Figure 5a**). Because our constructs carry aspartate substitutions in place of the two phosphoserines and adopt a closely similar binding mode to native phosphorylated tails, these phosphomimetics serve as a reliable model of the phosphorylated state (**S_Figure 5b**).

In addition to these differences, in the cryo-EM structures of the variant complexes, a Zn(II) ion is coordinated by two histidines of the SMAD4 MH2 domain located near the binding site of the SMAD3 C-terminus. In the I500V complex (Map 1), the coordination sphere formed by His382 and His405 (2.3 Å, AB interface) is completed by two ordered water molecules (2.1 Å) that contact the backbone carbonyl oxygens of Pro417 and Ile419 in the SMAD3 tail (**Figure 5b**). A similar feature is observed in the R496C complex. Because this cation-binding site has not been described in previous SMAD structures and was not detected in our WT cryo-EM complex (likely due to higher DTT concentrations during grid preparation), we searched for it in homotrimeric crystals of isolated wild-type SMAD4. In the 2.06 Å crystal structure that crystallized in the P36 SG, Zn(II) occupies the same site (within 1.2 Å) and is coordinated by the same two histidines in both chain B (2.05 and 1.93 Å) and chain A (2.00 and 2.10 Å). However, in the crystal lattice, the two water ligands seen in the cryo-EM structures are replaced by Cys345 and Asp360 from a symmetry-related molecule in chain B (2.24 and 1.80 Å), and by His371 from a symmetry mate (2.05 Å) together with a phosphate oxygen (2.34 Å) in chain A (**S_Figure 5c**).

#### 3.2 SMAD4 occupies the R-SMAD phosphotail pocket with an unphosphorylated C-terminus

SMAD4 is the only subunit whose C-terminus cannot be phosphorylated, yet it binds to the same pocket as the SMAD3 C-termini (**Figure 5a**). At the position equivalent to Ser425, SMAD4 carries Asp552, which is also its terminal residue. In the modeled orientation of the I500V cryo-EM complex (Map 1), where the C-terminus is fully traceable, both the side-chain and terminal carboxylate groups of Asp552 engage the first basic cluster of SMAD3, primarily contacting Lys378. The adjacent Leu551 fills this subsite hydrophobically, packing against Leu285, His289, and Val377 (**Figure 5b**). The C-terminal aspartate of SMAD4 thus acts as a constitutive, phosphorylation-independent mimic of the phosphorylated R-SMAD tail. This configuration also provides a structural rationale for the stimulatory effect of p300/CBP-mediated acetylation of SMAD3 Lys378 on SMAD-driven transcription <u>(Inoue et al. 2007)</u>. Lys378 lies alongside the SMAD4 tail, with its ε-amino group pointing toward the SMAD4 C-terminus. Acetylation extends this aliphatic surface into a subsite that is already hydrophobic in the SMAD4-bound complex, while the amide NH of acetyl-lysine retains the capacity to donate a hydrogen bond to the Asp552 carboxylate, accounting for the coactivator-driven enhancement of transcriptional output.

Taken together with the interfaces described above, these structures show that the two SMAD3 subunits are anchored to SMAD4 via distinct sets of interactions. Chain A engages SMAD4 through its C-terminal tail, which occupies the SMAD4 basic pocket and packs near the Zn(II) site, where Arg420 contributes to the local coordination environment in the I500V model. Conversely, Chain C engages SMAD4 through the β10/H4 surface harboring the Myhre positions and by receiving the SMAD4 C-terminus into its own basic pocket. Because His382 and His405 are unique to SMAD4 and absent in SMAD3, Zn(II) coordination is restricted to the SMAD4 pocket engaging Chain A, further distinguishing the two SMAD3 C-termini within the complex. The heterotrimer is thus assembled from two structurally non-equivalent SMAD4–SMAD3 interfaces, with the Myhre substitutions acting exclusively on one of them. Notably, the SMAD4 C-terminus is only partially resolved in the WT complex and untraceable in the R496C complex; whether these differences reflect intrinsic conformational flexibility, stabilization in I500V, or variations in sample preparation remains open.

#### 3.3 The two SMAD3 subunits adopt different N-terminal conformations

Although the SMAD3 segment preceding the MH2 domain (**S_Figure 1a**) adopts no regular secondary structure, three-dimensional variability analysis of the I500V heterotrimer followed by focused classification resolved a state (map 2) in which the two SMAD3 chains place this segment in clearly different positions (**Figure 5d**).

In chain A the segment folds back onto the domain from which it emerges. Tyr226 stacks against the indole of Trp406 in helix H5 (3.3 Å between ring atoms), Leu219 packs against Val356 (3.4 A), and the chain enters the domain through Phe231 and Trp232, which contact Ala250 and Asn401 (3.1-3.2 Å). Over residues 219 to 230 this segment makes no contact with the other SMAD3 subunit or with SMAD4, its only inter-subunit approach being Pro229 to SMAD4 Ser357 at 3.5 Å. The conformation is therefore intramolecular, and it is the better ordered of the two (mean B 118 Å^2^ against 159 Å^2^ for the equivalent segment of chain C).

In chain C the same segment instead extends across to the neighboring SMAD3 subunit. Gln222 is the anchor point, hydrogen bonding to Gln252 (2.9 Å) and Ser266 (3.5 Å) of chain A. Chain C could be traced from position 209, ten residues beyond the start of chain A, but the mean B factor of residues 209 to 218 exceeds 300 Å^2^ and we therefore assign no specific interactions in this stretch. Its significance is positional: the traced chain reaches Ser213, the most C-terminal residue of the regulatory cluster phosphorylated by GSK3 and CDK8/9 (Ser204, Ser208, Ser213; <u>(Aragón et al. 2011; Alarcón et al. 2009)</u>), placing this cluster immediately beyond the point at which the chain leaves the SMAD3-SMAD3 interface, while Ser204 and Ser208 remain disordered.

The two conformations have each been described previously, but only in binary complexes of SMAD2 or SMAD3 with a partner protein, where a cofactor contributes part of the binding surface (**Figure 5e**). Observing both conformations within a single cofactor-free heterotrimer indicates that they represent intrinsic conformational states of the R-SMAD rather than adaptations induced by partner binding, and that the two SMAD3 positions in the heterotrimer are not interchangeable with respect to this segment. These states are likely to be accessible to the WT complex as well; they were resolved in the I500V reconstruction, where variability analysis and focused classification were applied.

## Discussion

Our data show that Myhre syndrome substitutions at two SMAD4 positions converge on a single outcome: a longer-lived SMAD3–SMAD4 heterotrimer, driven by rewired polar contacts at a specific interface. The MH2 heterotrimer comprises three homologous yet non-identical interfaces—even between the two SMAD3 subunits (**Figure 3,5**). Because the Myhre positions lie in the β10/H4 area, the variants selectively affect the interface where SMAD4 serves as the contact donor. Rather than altering the overall MH2 fold or enlarging the buried surface area, these variants reorganize the polar contact network, forming hydrogen bonds in our crystal structures that are somewhat shorter than standard protein metrics <u>(Kemp et al. 2021; Cleland 2000)</u>. Although the affinity gain is modest in free-energy terms (2.0 to 4.6 kJ/mol across the five variants), it translates to a 4.7-fold increase in complex half-life. Notably, this stabilization is not an intrinsic property of the isolated MH2 domain—which is marginally less stable than WT—but emerges strictly upon heterotrimer assembly. A similar uncoupling of subunit versus complex stability occurs in platelet-type von Willebrand disease, where gain-of-function substitutions destabilize isolated monomers but increase target affinity upon binding <u>(Tischer et al. 2019)</u>. In a panel of SMAD4 variants drawn from cBioPortal, substitutions at these contact-forming residues modulate SMAD3 stabilization bidirectionally. The sharpest comparison occurs at Arg496: substitution to cysteine raises the SMAD3 melting temperature by 0.8 °C relative to WT, whereas substitution to histidine lowers it—a difference dictated by side-chain chemistry rather than position alone. Thus, the SMAD4 β10/H4–SMAD3 H1 interface acts as a tunable dial where substitutions stabilize or destabilize the complex depending on the specific contacts they add, modify, or disrupt. Beyond the core interface, our cryo-EM reconstructions resolve regulatory elements undetected in previous crystal structures. Specifically, we identified a Zn(II) cation coordinated by two SMAD4 histidines that chemically distinguishes the two SMAD3 tails, and we traced the unphosphorylated SMAD4 C-terminus—which is disordered in homotrimer crystal structures but ordered in our heterotrimer reconstructions, where it acts as a structural mimic of the phosphorylated R-SMAD tail.

Heterotrimer stabilization by Myhre syndrome variants also provides a structural framework for how inhibitory monoubiquitination regulates complex assembly. TRIM33-mediated monoubiquitination of SMAD4 at Lys519 <u>(Dupont et al. 2009)</u> and SMURF2-mediated modification of R-SMADs at Lys333 and Lys378 <u>(Tang et al. 2011; Zhou et al. 2017)</u> block subunit association, an effect reversed by USP9x and USP4. In our heterotrimer structures, all three lysines point directly into a subunit interface, lying within 3–4 Å of the adjacent protomer. Because a conjugated ubiquitin would sterically clash with the neighboring subunit, monoubiquitination and heterotrimer assembly are mutually exclusive, requiring free subunits as substrates (consistent with increased SMAD3–SMAD4 association upon TRIM33 depletion; <u>(Luo et al. 2026)</u>. By shifting the equilibrium toward assembled complexes, Myhre variants would reduce the pool of free, modifiable subunits—decreasing ubiquitination simply through reduced substrate availability rather than altered ligase activity (**Figure 6**).

**Figure 6.**
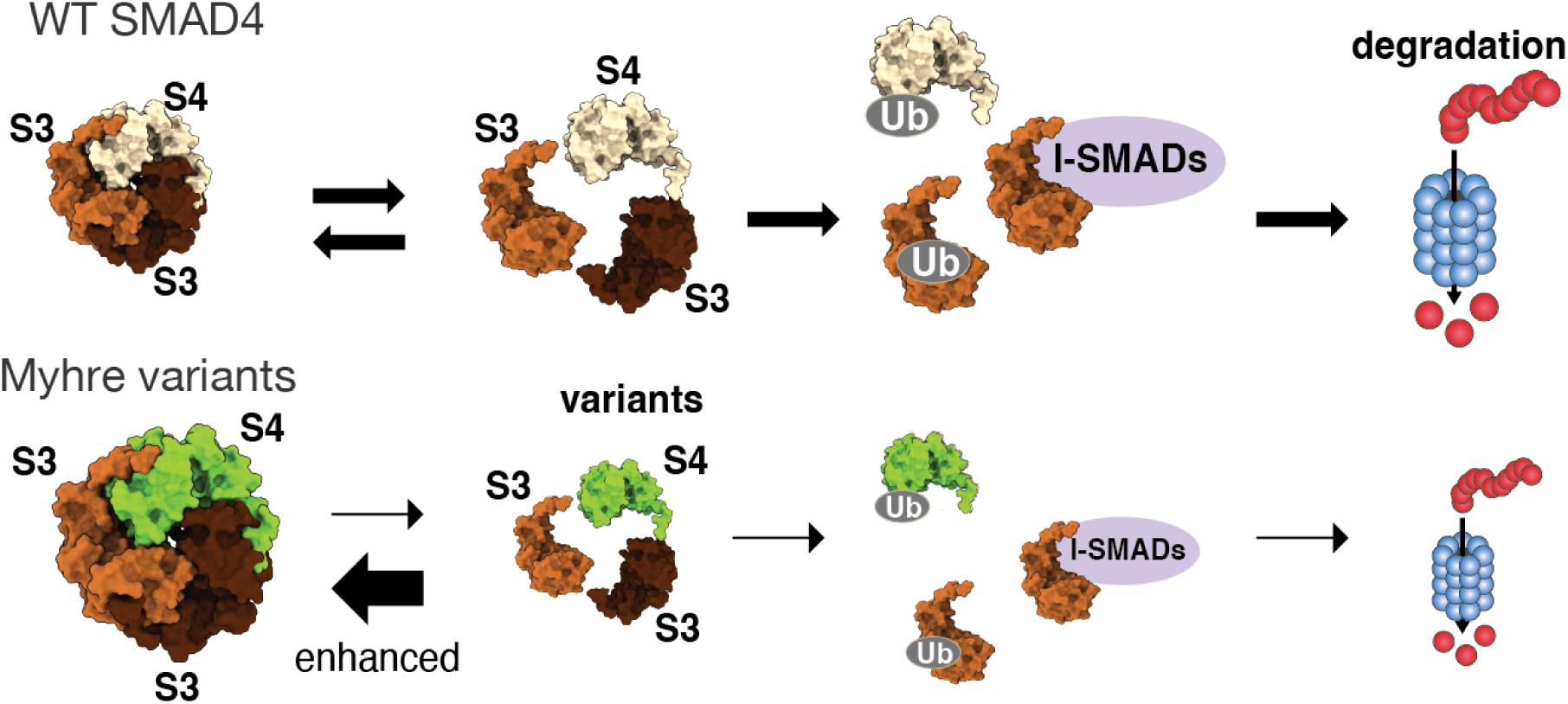
Working model. In the wild-type heterotrimer (top) assembly is reversible, and the dissociated subunits are ubiquitinated and displaced by inhibitory SMADs, leading to turnover. The Myhre variants (bottom) shift the equilibrium toward the assembled complex so that the subunits spend less time in the free state and ubiquitination and I-SMAD binding occur less often. S3, SMAD3; S4, SMAD4; Ub, ubiquitin.

The same reasoning extends to the inhibitory SMADs: SMAD6 and SMAD7 compete with SMAD4 for binding to activated R-SMADs <u>(Itoh et al. 2008; Miyazawa and Miyazono 2017; Hata et al. 1998)</u>, and the SMAD7 MH2 domain presents a basic groove - the same surface through which SMAD4 and the R-SMADs receive a phosphorylated C-terminal tail <u>(Murayama et al. 2020)</u>. Because each heterotrimer contains two SMAD3 subunits and the variant complexes release them 4.7-fold more slowly, the pool available to the inhibitory SMADs is reduced accordingly, again without any change in the affinity of the inhibitory SMAD itself (**Figure 6**).

The cellular consequences of these differences do not reduce to a simple gain in pathway output. While the initial description of Myhre syndrome linked impaired SMAD4 ubiquitination to reduced expression of TGF-β target genes <u>(Le Goff et al. 2011)</u>, patient-derived nasal epithelial cells carrying I500V, R496C, or I500T show elevated transcriptional activity attributed to tighter SMAD complex assembly <u>(Lindsay et al. 2025)</u>, an interaction our measurements confirm is both stronger and longer-lived. Yet in those same cells, heightened signaling is accompanied by reduced proliferation, impaired differentiation, and defective bacterial clearance, showing that stronger signaling does not translate into better cellular function. Our data suggest a mechanism to reconcile these findings: prolonging complex lifetime would sustain promoter occupancy while reducing pathway turnover and the number of completed signaling cycles. The net functional outcome of these opposing effects would therefore depend on promoter context and stimulus duration. Because the primary consequence of Myhre substitutions is the persistence of assembled complexes rather than the formation of a dominant toxic species, strict allele selectivity may not be required to achieve therapeutic benefit <u>(De Falco et al. 2025)</u>. Compounds that accelerate heterotrimer dissociation—or restore normal complex turnover by other means—could therefore hold therapeutic potential, even if they also act on the wild-type complex. Collectively, five substitutions at two SMAD4 positions prolong the lifetime of the SMAD3–SMAD4 heterotrimer by rewiring polar contacts at the very same interface where other mutations impair complex stability. The atomic-resolution structures reported here define the contact network underlying this kinetic stabilization and provide a molecular framework for the rational discovery of modulators targeting SMAD complex turnover in Myhre syndrome.

## Supplementary Tables

**Supplementary Table 1:** Abridged data collection and refinement statistics for the non-deposited X-ray structures compared in this work. Data collection and refinement statistics for the structures determined in this work. All of them crystallized in space group F4_1_32 with one molecule in the asymmetric unit and were refined at resolution of 1.74-2.13 Å. Residues and mean B factor refer to the modelled protein of the chain used for analysis. Variants are named by the SMAD4 substitution they carry; the wild-type residues at the two positions, Arg496 and Ile500, are present in every structure except where substituted.

| Variant | Crystal | Space group | Cell (Å) | Resol. (Å) | Rwork | Rfree | Resid. | Mean B (Å<sup>2</sup>) |
|---|---|---|---|---|---|---|---|---|
| I500V | I500V (II) | F4132 | 196.2 | 1.74 | 0.1742 | 0.2139 | 193 | 46.4 |
| I500V | I500V (III) | F4132 | 196.6 | 1.95 | 0.1956 | 0.2294 | 197 | 29.7 |
| I500M | I500M (II) | F4132 | 196.3 | 1.88 | 0.1789 | 0.2205 | 192 | 60.1 |
| R496C | R496C (II) | F4132 | 196.8 | 1.76 | 0.1556 | 0.1935 | 194 | 51.4 |
| R496C | R496C (III) | F4132 | 196.1 | 1.96 | 0.1883 | 0.2296 | 190 | 32.3 |
| R496C | R496C (IV) | F4132 | 196.3 | 2.13 | 0.215 | 0.2533 | 190 | 35.4 |

**Supplementary Table 2:**
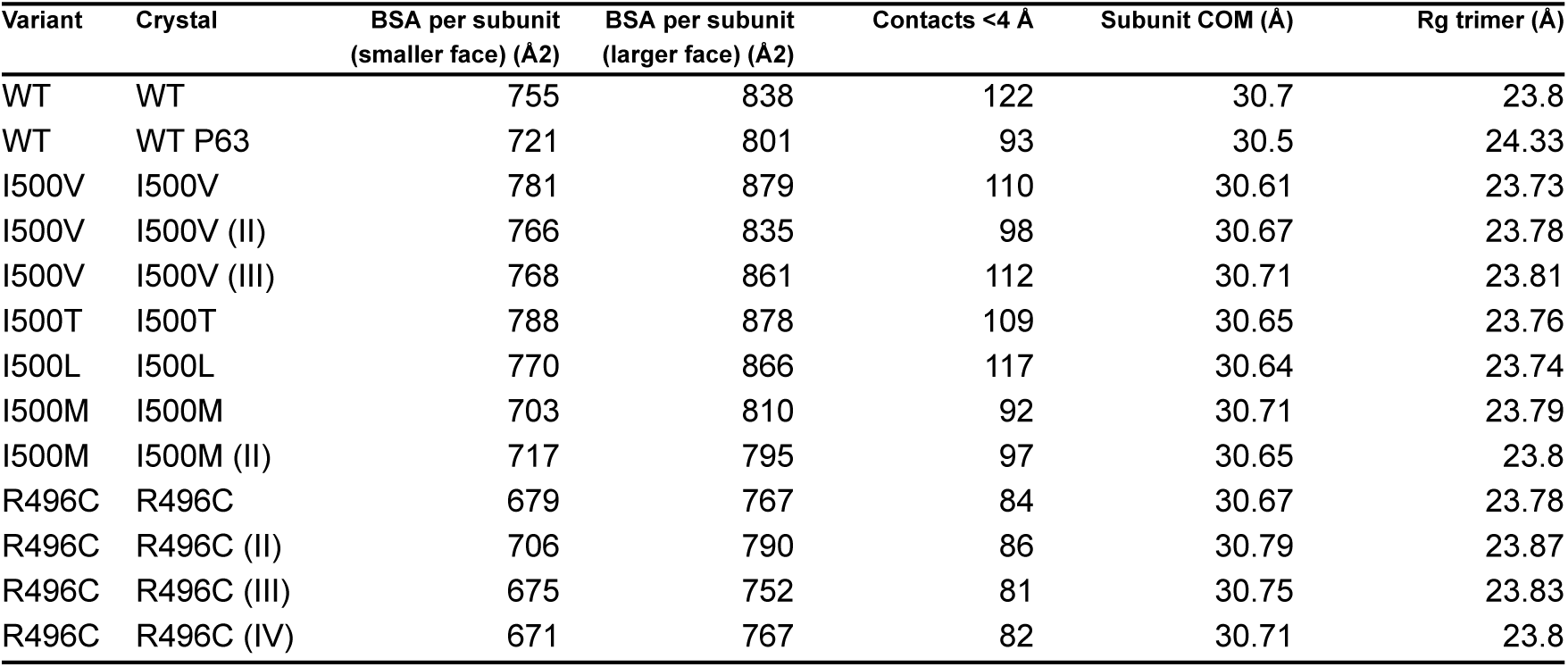
Interfacial surface area and geometry. Interface and global dimensions computed on the homotrimer generated by the crystallographic three-fold. To make the thirteen structures directly comparable, all values use the residue set common to every structure (173 residues, 320-542); absolute values therefore differ from those obtained on the full modelled chain, while the comparison between structures is unaffected. Solvent-accessible surface was computed by the Shrake-Rupley method with a 1.4 Å probe and 128 sphere points, excluding hydrogens and using the first alternate conformation where present. Buried surface area (BSA) per subunit is the loss of solvent-accessible surface of one subunit on forming the interface with its neighbour; contacts are heavy-atom pairs within 4 Å across the same interface; subunit COM is the mean separation of the three subunit centres of mass; Rg is the radius of gyration of the trimer. Subunit separation spans 30.58-30.90 Å and the radius of gyration 23.71-23.92 Å across all thirteen structures, so the variants are not more compact than the wild type. Buried surface and contact number straddle the wild-type values in both directions: I500V, I500T and I500L bury more surface than either wild-type crystal and I500M and R496C bury less.

| Variant | Crystal | BSA per subunit (smaller face) (Å2) | BSA per subunit (larger face) (Å2) | Contacts <4 Å | Subunit COM (Å) | Rg trimer (Å) |
|---|---|---|---|---|---|---|
| WT | WT | 755 | 838 | 122 | 30.7 | 23.8 |
| WT | WT P63 | 721 | 801 | 93 | 30.5 | 24.33 |
| I500V | I500V | 781 | 879 | 110 | 30.61 | 23.73 |
| I500V | I500V (II) | 766 | 835 | 98 | 30.67 | 23.78 |
| I500V | I500V (III) | 768 | 861 | 112 | 30.71 | 23.81 |
| I500T | I500T | 788 | 878 | 109 | 30.65 | 23.76 |
| I500L | I500L | 770 | 866 | 117 | 30.64 | 23.74 |
| I500M | I500M | 703 | 810 | 92 | 30.71 | 23.79 |
| I500M | I500M (II) | 717 | 795 | 97 | 30.65 | 23.8 |
| R496C | R496C | 679 | 767 | 84 | 30.67 | 23.78 |
| R496C | R496C (II) | 706 | 790 | 86 | 30.79 | 23.87 |
| R496C | R496C (III) | 675 | 752 | 81 | 30.75 | 23.83 |
| R496C | R496C (IV) | 671 | 767 | 82 | 30.71 | 23.8 |

**Supplementary Table 3:** Expression constructs used in this study. For each construct the table provides the expression vector together with the protease used to remove the tag. Molecular weights are calculated for the tag-free protein, except for SUMO SMAD4 MH2_314, where the value includes the His–SUMO tag. The full-length SMAD4 construct was used uncleaved. Superscript letters identify the source of each vector: a, <u>(Berrow et al. 2007)</u>; b, <u>(Dümmler et al. 2005)</u>; c, <u>(Scholz et al. 2013)</u>.

| Protein | Tag | M.W. (kDa) | Notes |
|---|---|---|---|
| SUMO SMAD4 MH2_314<br>(314–552) | His-SUMO tag | 38 | pOPINS<sup>a</sup>; cleavable by SUMO protease<br>Uniprot Q13485 |
| SMAD4 MH2_314<br>(314–552) | His | 26 | pETM11<sup>b</sup>; tag cleaved by TEV protease.<br>Uniprot Q13485 |
| SMAD4 MH2_272<br>(272–552) | His tag | 31 | pETM11<sup>b</sup>; tag cleaved by TEV protease<br>Uniprot Q13485 |
| Full-length SMAD4 (1-552) | Twin-Strep tag | 64 | pCoofy34<sup>c</sup>; Twin-Strep-tag and a 3C protease cleavage site, uncleaved Uniprot Q13485 |
| SMAD1 MH2_EEE<br>(259-465) | His-SUMO tag | 24 | pOPINF<sup>a</sup>; His-tag cleaved by 3C<br>Uniprot Q15797 |
| SMAD2 MH2_DMD<br>(231-467) | His-SUMO tag | 27 | pOPINF<sup>a</sup>; His-tag cleaved by 3C<br>Uniprot Q15796 |
| SMAD3 MH2_DVD<br>(189-425) | His-SUMO tag | 27 | pOPINF<sup>a</sup>; His-tag cleaved by 3C<br>Uniprot P84022 |

**Supplementary Figure 1.**
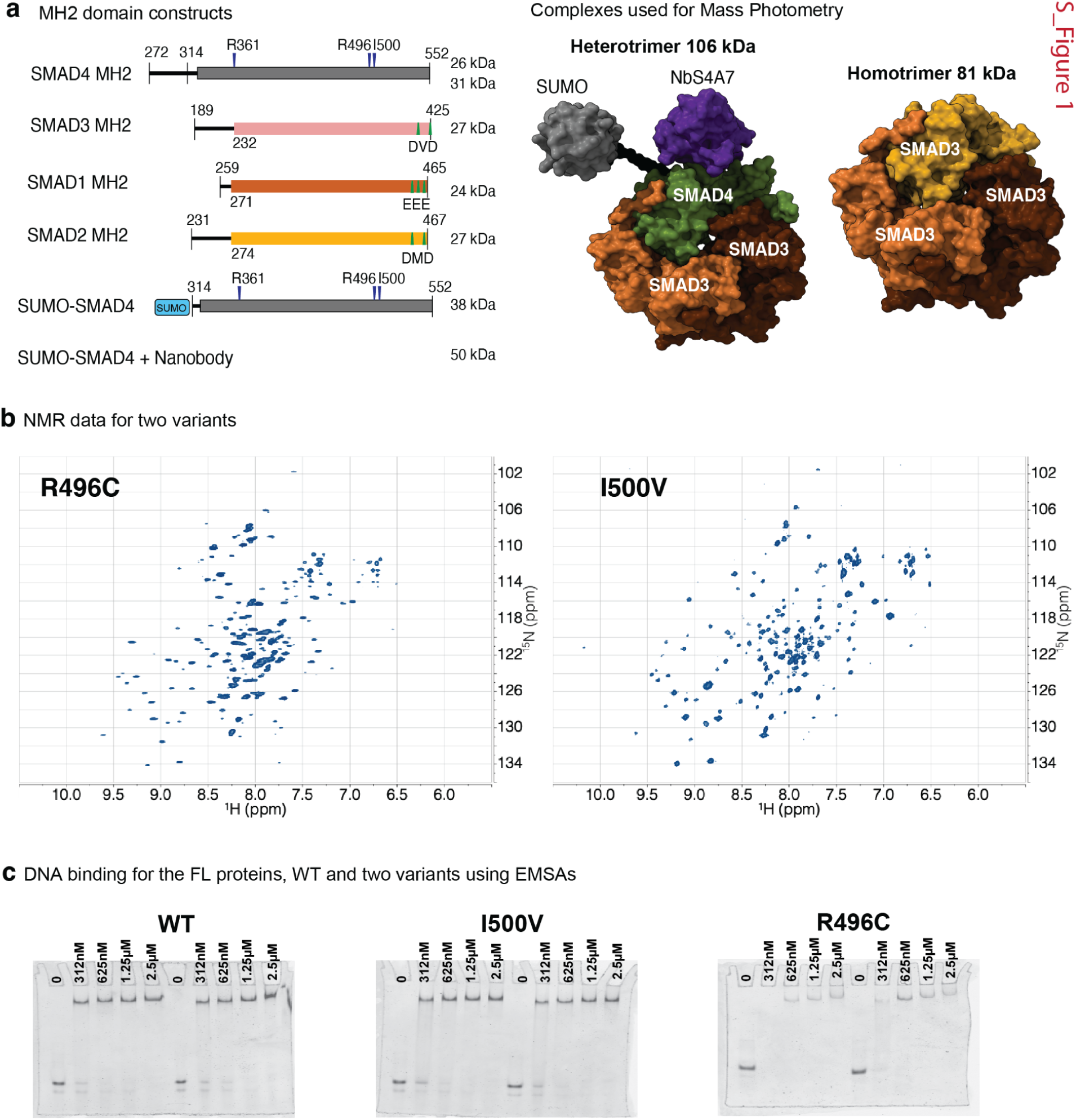
Construct design, structural assemblies for mass photometry, and biochemical characterization. **(a)** Schematic representation of MH2 domain constructs used in this study (left), highlighting residue boundaries, molecular weights, and positions of the MYHRS variants (Arg496 and Ile500) and of Arg361. Right panel displays surface representations of molecular complexes analyzed by Mass Photometry, including the SUMO–SMAD4:SMAD3:NbS4A7 heterotrimer (∼106 kDa) and the SMAD3 homotrimer (∼81 kDa). Plasmids are indicated in **S_Table 3**. **(b)** 1H–15N HSQC NMR spectra for I500V and R496C variants (600 MHz). **(c)** Electrophoretic Mobility Shift Assays (EMSAs) showing that the DNA binding capacity is retained (CY5-AAGATTGTGGATTGAGAATTCGGCAGCCCCGGCGCGGCCCCATAT) of full-length (FL) WT SMAD4, I500V, and R496C variants across increasing protein concentrations as indicated in the figure. DNA concentration is 7.5 nM.

**S_Figure 2.**
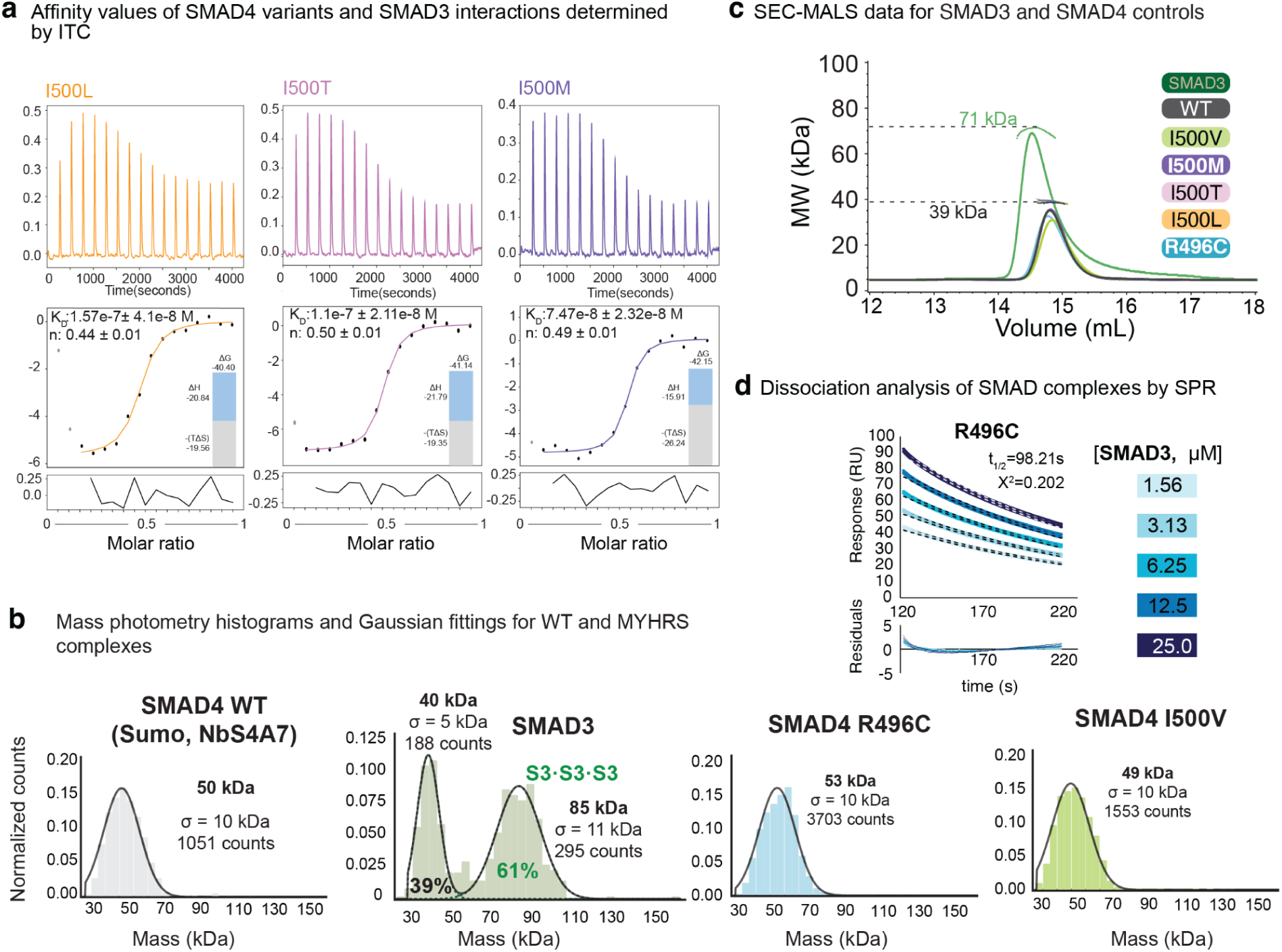
Additional SPR, mass photometry and SEC-MALS data. (a) Isothermal titration calorimetry (ITC) of SMAD3 binding to I500L (orange), I500T (orchid) and I500M (violet) measured at 37 °C. Top, raw thermograms. Middle, integrated heats fitted to a one-site binding model, with the fitted K_D_ and stoichiometry (n) shown in each panel. Insets show the thermodynamic signature (ΔG, ΔH, and −TΔS). Bottom, fit residuals. Grey points were excluded from the fit. (b) Mass photometry of the individual components: SMAD4 MH2 WT, R496C and I500V, and SMAD3 MH2, with Gaussian fits. Each population is labeled with its mean mass, σ, counts and fraction of total. SMAD3 alone forms several species including homotrimers (S3·S3·S3). (c) SEC-MALS controls of SMAD3 MH2 alone and of SMAD4 MH2 WT and variants alone. Dashed lines mark 71, 53 and 39 kDa. (d) SPR dissociation phases for SMAD3 MH2 (1.56–25 µM) from immobilized SMAD4 MH2 R496C. Colored lines are data and dashed black lines are fits, with residuals below. t_1/2_ and χ² are indicated.

**Supplementary Figure 3.**
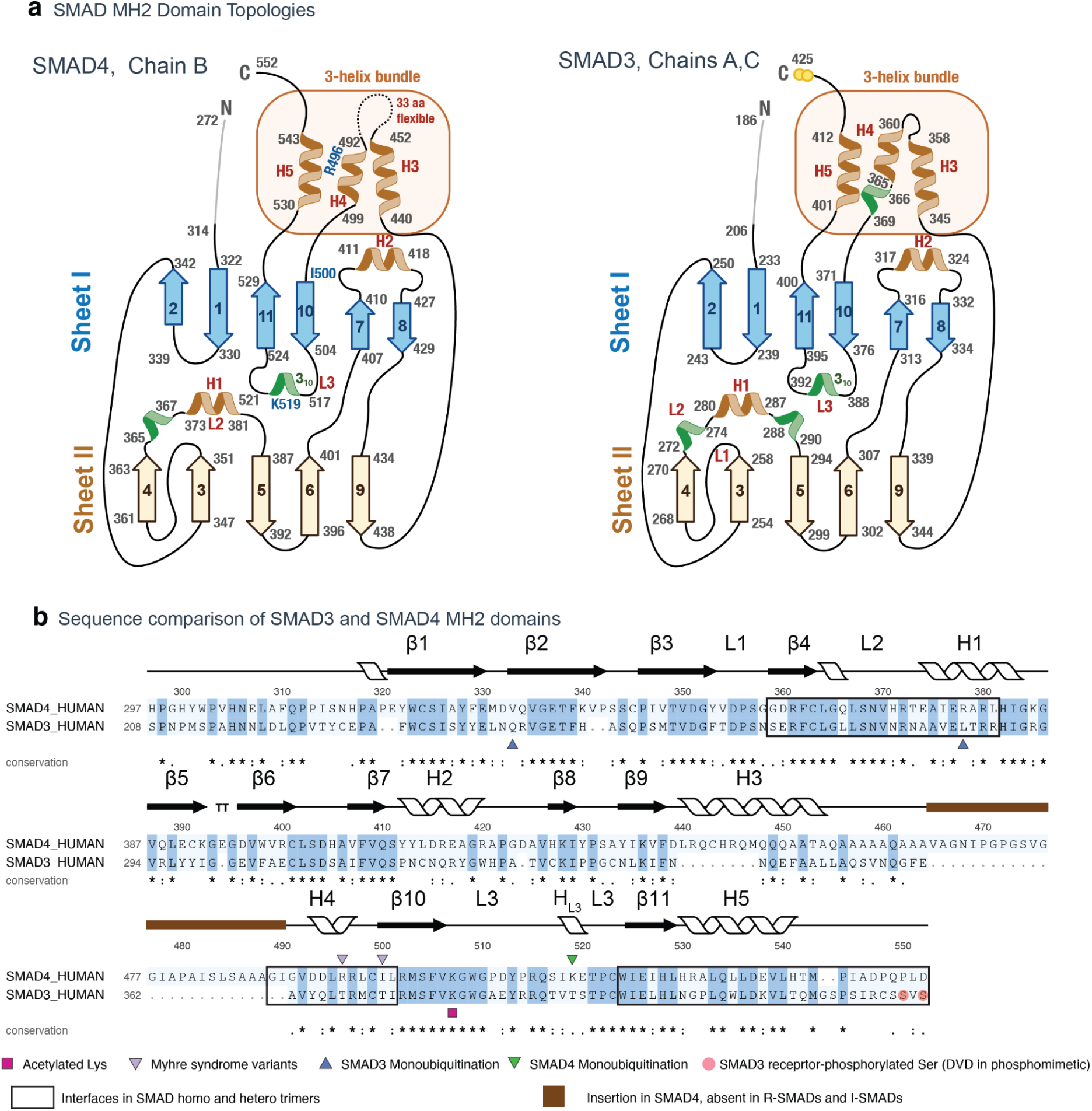
Topology and sequence comparison of the SMAD4 and SMAD3 MH2 domains. (c) Topology diagrams of the MH2 domains of SMAD4 (chain B) and SMAD3 (chains A, C), showing β-sheets I and II, helices H1–H5, the three-helix bundle and loops L1–L3. The flexible 33-residue SMAD4 insertion, the Myhre syndrome residues R496 and I500, and K519 are indicated. (d) Structure-based alignment of the SMAD4 and SMAD3 MH2 domains with secondary structure shown above. Boxes mark interfaces in SMAD homo- and heterotrimers, and brown shading marks the insertion in SMAD4 that is absent from R-SMADs and I-SMADs. Purple triangles mark the Myhre syndrome variant positions. Green triangle, SMAD4 K519, monoubiquitinated by TRIM33. Blue triangles, SMAD3 K333 and K378, monoubiquitinated by SMURF2. Magenta square, acetylated K378. Red circles, receptor-phosphorylated SMAD3 serines (DVD in the phosphomimetic).

**Supplementary Figure 4.**
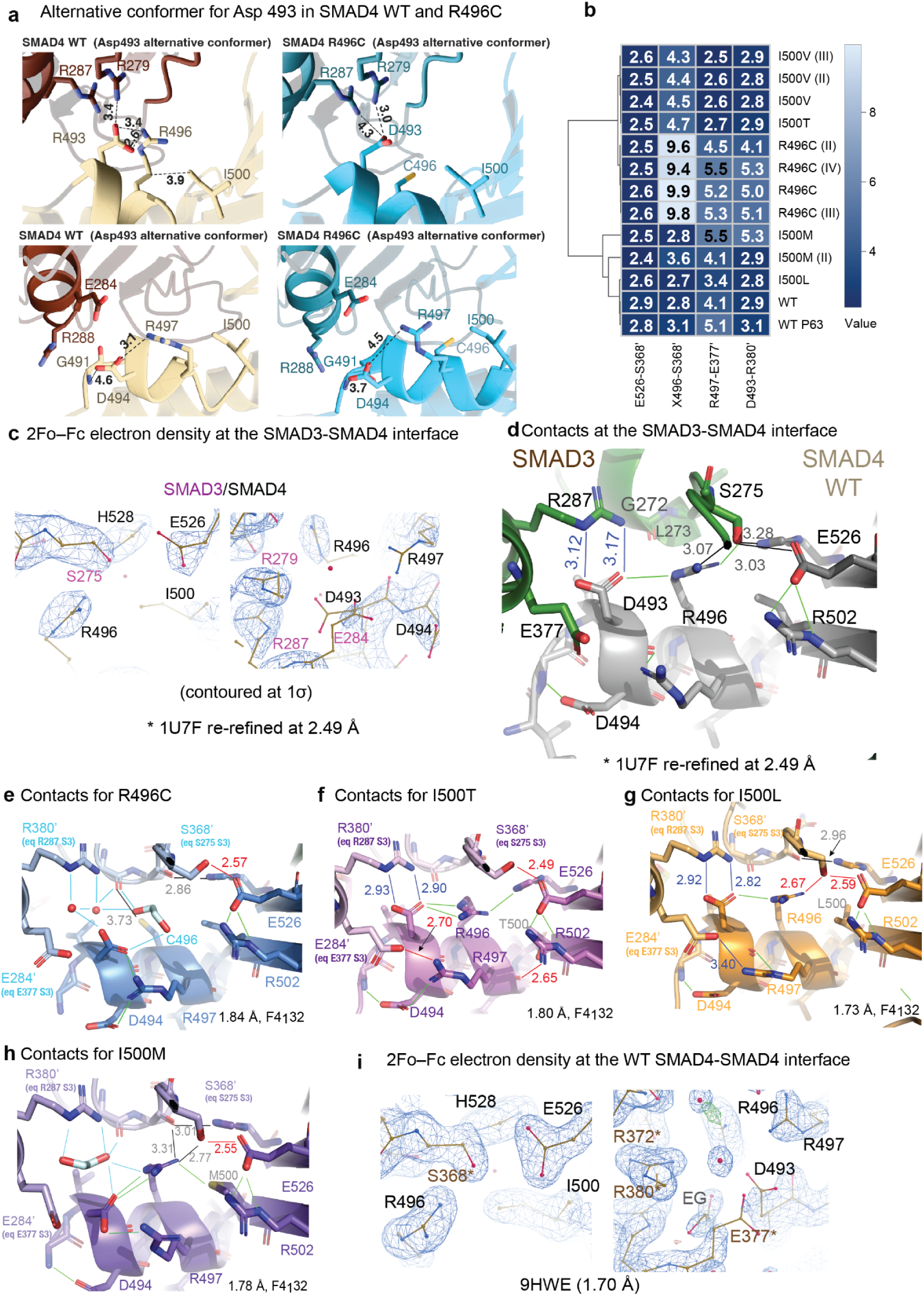
Inter-subunit contacts across the crystal series. (a) Alternative conformers for Asp493 in SMAD4 WT and R496C variant complexes. (b) Heatmap of the four inter-subunit distances measured in each of the thirteen crystal structures, clustered by Pearson distance. Columns are the Glu526-Ser368’, residue 496-Ser368’, Arg497-Glu377’ and Asp493-Arg380’ contacts; rows are individual crystals, with replicates of the same variant numbered in parentheses. Residue 496 is measured from the guanidinium in arginine-containing structures and from the thiol in R496C. (c) 2Fo–Fc electron density (contoured at 1σ) at the SMAD3-SMAD4 interface of PDB 1U7F, re-refined at 2.49 Å. SMAD3 residues are labelled in magenta and SMAD4 residues in black. (d) The same interface in the re-refined 1U7F structure, with SMAD3 in green and SMAD4 in grey. Distances in angstroms; line colors as in Figure 4d. (e-h) The equivalent region in the crystal structures of R496C (e), I500T (f), I500L (g) and I500M (h), each with its resolution and space group indicated. Primed residues belong to the symmetry-related subunit that completes the homotrimer; equivalent SMAD3 residues are given in parentheses where shown. (i) 2Fo–Fc electron density (contured at 1σ) for the wild-type SMAD4 crystal structure over the same residues. Asterisks mark residues from the neighbouring subunit.

**Supplementary Figure 5.**
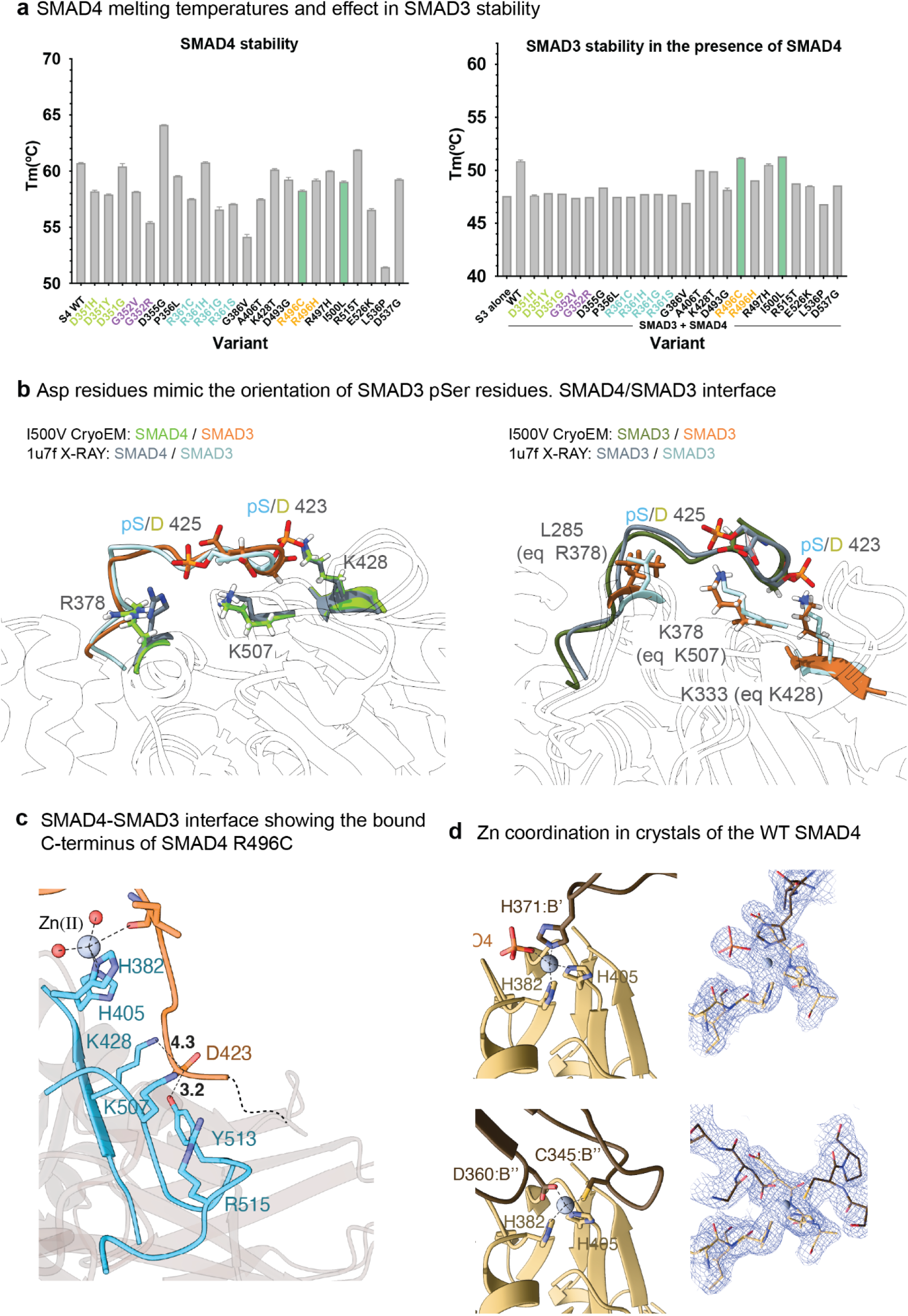
Phosphomimetic Asp residues adopt the conformation of SMAD3 phosphoserines, and Zn(II) coordination in WT SMAD4 crystals. (a) Tm of SMAD4 variants alone (left) and of SMAD3 mixed with each SMAD4 variant (right), measured by nanoDSF; dotted lines mark the T_m_ of WT SMAD4 and of SMAD3 alone. Bars show mean ± s.d.; n = 4 technical replicates. (b) Superposition of the I500V cryo-EM structure on the crystal structure of the phosphorylated SMAD3–SMAD4 complex (PDB 1U7F). Left, SMAD4– SMAD3 interface: SMAD4 in green (cryo-EM) or grey (1U7F) and SMAD3 in orange (cryo-EM) or pale cyan (1U7F). The phosphomimetic SMAD3 D423 and D425 overlay pS423 and pS425 of 1U7F and contact SMAD4 R378, K428 and K507. Right, SMAD3–SMAD3 interface: SMAD3 subunits in dark green and orange (cryo-EM) or grey and pale cyan (1U7F). The C-terminal residues contact the equivalent SMAD3 residues L285, K333 and K378, which correspond to SMAD4 R378, K428 and K507. (c) Zn(II) coordination in crystals of WT SMAD4 MH2. Top, a Zn(II) ion coordinated by H382 and H405 of chain A, H371 of a symmetry-related chain (B′) and a phosphate ion. Bottom, a second Zn(II) site formed by H382 and H405 of chain B and D360 and C345 of a symmetry-related chain (B″). Left, stick models with coordination bonds (green dashed lines). Right, 2Fo–Fc electron density (contoured at 1σ). Primes denote symmetry-related molecules. The Zn(II) site formed by SMAD4 H382 and H405 in the cryo-EM structure is shown in Figure 5b.

**Schematics 1.**
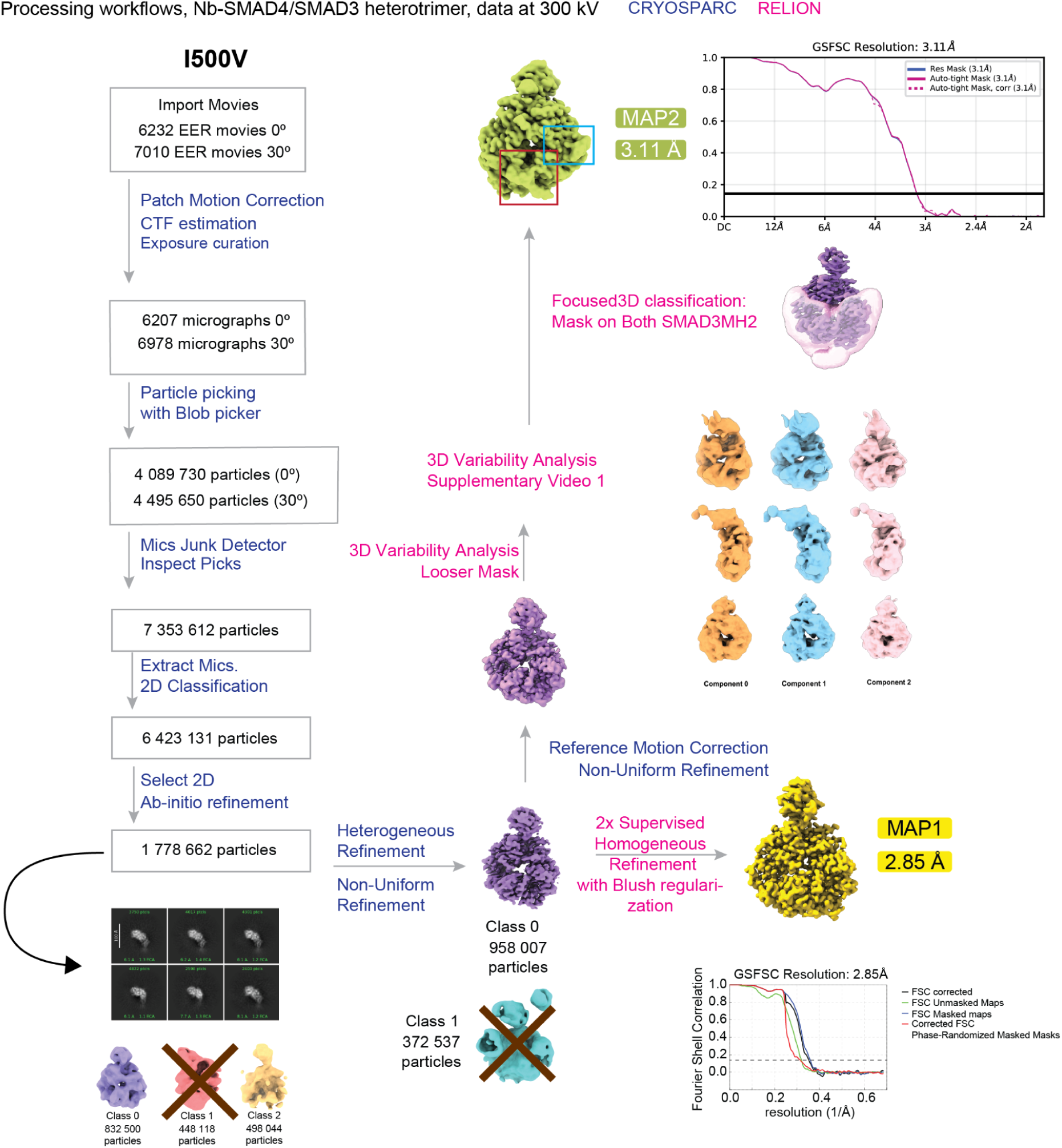
Cryo-EM processing workflow for the NbS4A7–SMAD4 I500V–SMAD3 complex. Data were collected at 300 kV at 0° and 30° stage tilt. Steps in blue were done in cryoSPARC and steps in pink in RELION. Representative 2D class averages are shown (scale bar, 100 Å). After heterogeneous refinement, class 0 (958,007 particles) was refined to MAP1 (2.85 Å). Focused 3D classification with a mask on both SMAD3 MH2 domains gave MAP2 (3.11 Å; I500V class 2). Insets show gold-standard FSC curves; dashed lines indicate FSC = 0.143. 3D variability analysis is shown in **Supplementary Video 1**.

**Schematics 2.**
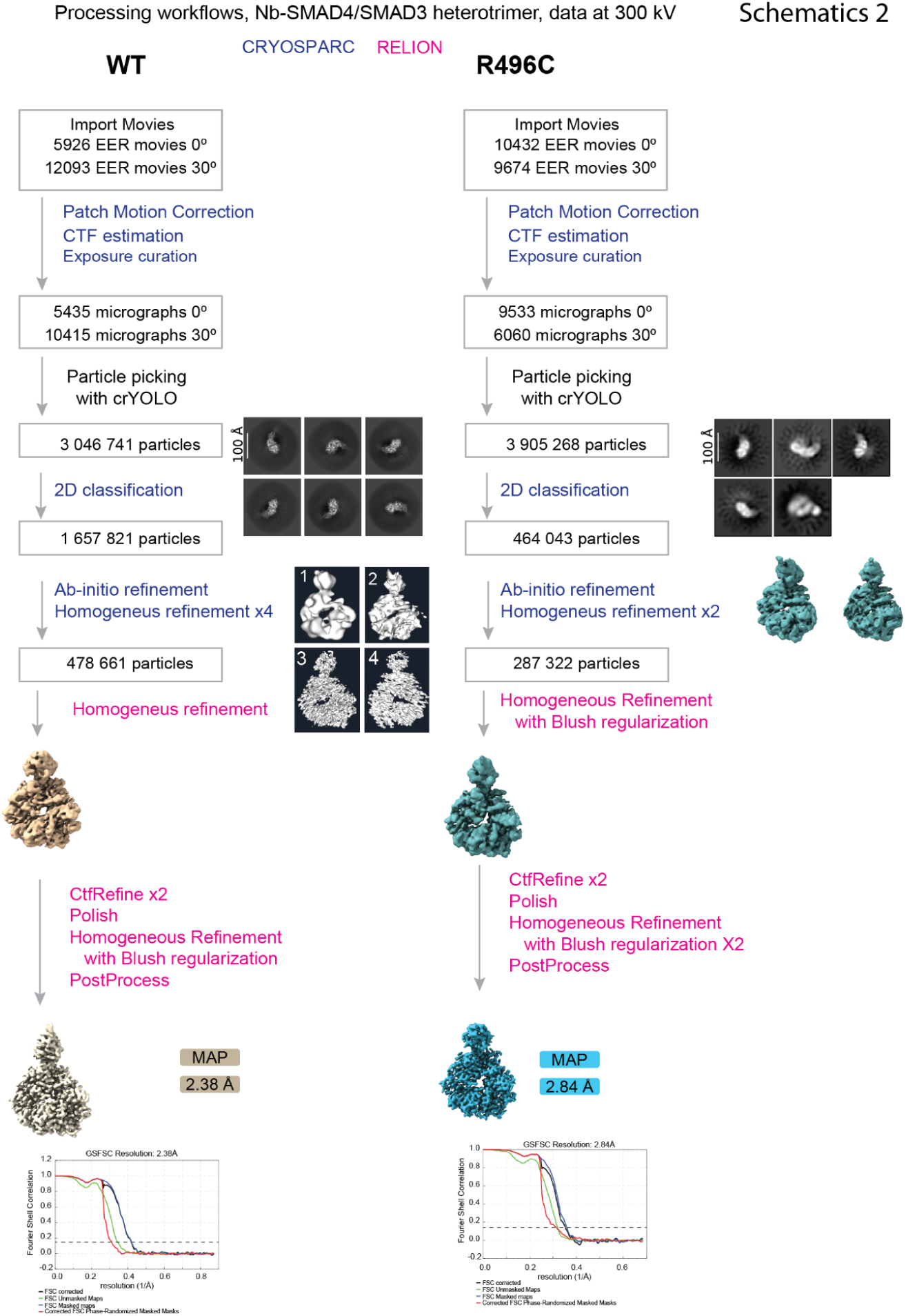
Cryo-EM processing workflows for the NbS4A7–SMAD4 WT–SMAD3 and NbS4A7–SMAD4 R496C–SMAD3 complexes. Data were collected at 300 kV at 0° and 30° stage tilt. Steps in blue were done in cryoSPARC and steps in pink in RELION. Particles were picked with crYOLO. Representative 2D class averages are shown (scale bars, 100 Å). Final maps reached 2.38 Å (WT) and 2.84 Å (R496C). Bottom, gold-standard FSC curves (corrected, unmasked, masked and phase-randomized); dashed lines indicate FSC = 0.143

## Methods

### Protein cloning, expression, and purification

Codon-optimized constructs corresponding to human SMAD proteins (SMAD4 MH2_314, extended SMAD4 MH2_272, full-length SMAD4, and phospho-mimetic SMAD1-EEE and SMAD2 and SMAD3-DVD MH2 domains) were cloned into vectors specified in **S_Table 3** and in the literature <u>(Condeminas et al. 2026; Macias et al. 2025)</u>. Recombinant SMAD proteins were expressed in *Escherichia coli* B834(DE3) in LB-TB media to provide extra amino acids and vitamins as this strain cannot make its own methionine, and supplemented with glycerol and appropriate antibiotics. For ^15^N labeled proteins we used BL21 cells and M9 minimal medium supplemented with ^15^N-ammonium chloride as the sole nitrogen source. Pre-cultures were grown in unlabeled M9 to adapt cells to minimal media. After a pellet-wash step to avoid unlabeled background minimal media we inoculated 1 L of ^15^N-M9 minimal medium with 10–20 mL of overnight preculture (starting OD600 ≈ 0.05–0.1). Expression was induced with 0.5 mM IPTG at 0.8 OD600 followed by overnight incubation at 20 °C. Harvested cells were lysed enzymatically (lysozyme and DNase I) and then mechanically (EmulsiFlex-C3, Avestin, Ottawa, ON, Canada). His-tagged or SUMO-fused SMAD variants were isolated from the soluble fraction using a 5 mL HisTrap HP column (Cytiva, Marlborough, MA, USA) with an imidazole gradient (40–400 mM) using an AKTA FPLC (GE Healthcare, Chicago, IL, USA) or NGC Quest 10 PlusChromatography System (BIO-RAD, Hercules, CA, USA). Following cleavage of the His-SUMO tag by SUMO protease and subsequent removal of the cleaved tag via a second HisTrap step, proteins were further purified by size-exclusion chromatography (SEC) on a Superdex 75 HiLoad 16/600 column (Cytiva) in 20 mM Tris pH 7.5, 100 mM NaCl, 2 mM TCEP. Full-length SMAD4 constructs were purified using a StrepTrap column (Cytiva) eluted with 2.5 mM desthiobiotin prior to SEC. Proteins were eluted in20 mM Tris pH 7.5, 200 mM NaCl, 1 mM DTT. Purified proteins were concentrated to 15-20 mg/mL (SMAD4 MH2 domain WT and variants 272-552), 18–25 mg/mL (SMAD4 MH2 domain WT and variants 314-552) and ∼ 3-4 mg/mL for the FL SMAD4, SMAD1 and SMAD3 MH2 domains. All final protein preparations were verified for purity by SDS-PAGE and Mass Spectrometry, concentrated, flash-frozen in liquid nitrogen, and stored at −80 °C.

### NMR

NMR spectroscopy. Uniformly ¹⁵N-labelled SMAD4 MH2 domains (residues 314–552; WT, I500V and R496C) were expressed in E. coli BL21 in M9 minimal medium containing ¹⁵N-ammonium chloride as described above and purified as for the unlabelled proteins. Samples were prepared at 0.8 mM in 50mN NaPO_4_ and 100mM NaCl pH 5.0, supplemented with 10% (v/v) D₂O for the lock, in 3 mm tubes (total volume 160 µL). Two-dimensional ¹H–¹⁵N HSQC spectra of the WT domain were recorded at 298 K on a Bruker Avance 950 MHz spectrometer (Bruker BioSpin GmbH, Ettlingen, Germany) equipped with a cryogenic TCI probe at the IBS in Grenoble, France; spectra of the I500V and R496C variants were recorded at 298 K on a Bruker Avance 600 MHz spectrometer equipped with a cryogenic TCI probe. ¹H–¹⁵N HSQC spectra were acquired with spectral widths of 12 ppm (¹H) and 38 ppm (¹⁵N), 2048 × 256 complex points and 32 scans per increment, giving acquisition times of 1 h per spectrum. Spectra were processed with TopSpin 4. The spectra were used only to establish that the variants retain the folded WT architecture.

### Differential scanning fluorimetry (DSF) and nanoDSF

Initial experiments were performed using a Prometheus NT.48 system (NanoTemper, Munich, Germany) at the ProLinC Facility, University of Linköping, followed by final measurements on a Prometheus Panta (NanoTemper) system at the SPC facility, EMBL Hamburg. Protein samples were loaded into Prometheus High Sensitivity Capillaries (NanoTemper) and exposed to a temperature gradient (25–80 °C) at a rate of 1 °C/min. Excitation power was automatically determined using the "Discovery Scan" mode. For the analyses of cancer variants, proteins were diluted in 25 mM Tris pH 9.0, 150 mM NaCl, 1 mM TCEP, and 5% glycerol to minimize aggregation of the less stable mutants, whereas R-SMAD proteins were stored in 20 mM Tris pH 7.5, 100 mM NaCl, and 2 mM TCEP. For the experiment, volumes of protein complexes were adjusted to ensure identical final buffer compositions across all samples. Final assay measurements were performed at protein concentrations of 14.5 µM for the R-SMADs and 29 µM for the SMAD4 variants.

### Homogeneous Time-Resolved Fluorescence HTRF

HTRF was employed to evaluate alterations in SMAD4–R-SMAD heterotrimer complex formation using an indirect competition assay. The assay mixture comprised 100 nM His-SUMO-SMAD3 (residues 189–425, DVD construct) and 50 nM TwinStrepTag-SMAD4 MH2 domain (residues 314–552). SMAD4–SMAD3 complex formation was detected using an anti-6His-d2 monoclonal antibody (Revvity, Cat. No. 61HISDLA, Waltham, MA, USA) and Strep-Tactin®-Terbium Cryptate (Revvity, Cat. No. 610STTLA). Fluorescent conjugate dilutions and protein concentrations were optimized according to Revvity’s standard assay development guidelines. All reactions were conducted in HTRF PPI-Terbium Detection Buffer (Revvity, Cat. No. 61DB10RDF), which substantially improved the signal-to-noise ratio and signal stability over time compared to other evaluated buffers. For competitive binding experiments, untagged SMAD4 MH2 domain (residues 314–552) constructs—including wild-type (WT) and all reported Myhre syndrome (MYHRS) variants—were utilized as cold competitors. Prior to the assay, untagged variants underwent buffer exchange into HTRF protein buffer (20 mM Tris pH 7.5, 100 mM NaCl, 0.01% v/v Tween-20) using Zeba Spin Desalting Columns (7K MWCO, 30–130 µL; Thermo Fisher Scientific, Cat. No. 89882, Waltham, MA, USA). Serial dilutions of the competitor variants were dispensed into 384-well white ProxiPlates (Revvity) using an Echo 650 Liquid Handler (Beckman Coulter, Brea, CA, USA), ensuring an equivalent final volume of HTRF protein buffer across all wells. Tagged proteins (SMAD3 and SMAD4) and HTRF detection reagents (anti-6His-d2 and Strep-Tactin®-Tb) were prepared in HTRF PPI-Terbium Detection Buffer and dispensed sequentially using a Multidrop Combi Reagent Dispenser (Thermo Fisher Scientific) to reach a final reaction volume of 10 µL per well. Following a 30 min incubation at room temperature, time-resolved fluorescence was measured on an EnVision plate reader (Revvity) using a TRF excitation filter at 340 nm and emission filters at 620 nm (donor, Terbium) and 665 nm (acceptor, d2). To mitigate intra-plate signal drift, high and low control wells were included across each row. The HTRF ratio (665nm/620 nm x10^4^) was calculated, and percentage inhibition was derived relative to the control wells according to Equation 1. Half-maximal inhibitory concentration IC50 values were determined individually for each dose-response curve using GraphPad Prism 8 (GraphPad Software, Boston, MA, USA).

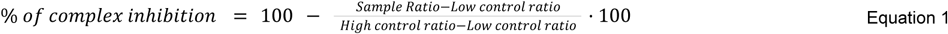

### Complex formation analysis using SEC-MALS

These experiments were performed using a modular HPLC Prominence system (Shimadzu, Kyoto, Japan) connected to an autosampler SIL-20AC, an LCD20-ADsp pump, an SPD-20A UV detector and two light-scattering detectors, a Dawn Heleos II (Wyatt, Santa Barbara, CA, USA) MALS and an Optilab t-Rex RI (Wyatt). Samples were prepared from frozen protein stocks. Proteins were thawed on ice and centrifuged at 17000*g* and 4 °C for 10 min. They were then filtered with 0.22 µm membrane filters and transferred to the SIL-20AC autosampler and injected in a Superdex 200 10/300 Increase column (Cytiva). Proteins were combined in a 2:1 ratio (SMAD4:SMAD3) and 130 µL or 190 µL was injected using 50 µM:25 µM and 100 µM:50 µM ratios. The running buffer was 20 mM Tris pH 7.5, 100 mM NaCl, 2 mM TCEP. Results were analyzed using ASTRA (Wyatt) and a Zimm model to calculate average MW of the selected fractions.

### Mass Photometry assays

Mass photometry (MP) experiments were performed at nanomolar protein concentrations using Refeyn OneMP and TwoMP instruments (Refeyn Ltd., Oxford, UK). All assay buffers (20 mM Tris pH 7.5, 100 mM NaCl) were filtered using Steriflip® Filter Units (0.22 µm, Merck KGaA, Darmstadt, Germany) and Amicon® Ultra 0.5 mL 10 kDa centrifugal filters (Millipore, Burlington, MA, USA) prior to use. For each acquisition, minimal background noise was confirmed by measuring a buffer control drop prior to sample addition.

For OneMP acquisitions, mass calibration was performed using NativeMark™ Unstained Protein Standard (Life Technologies, Cat. No. LC0725; 66 kDa, 146 kDa, 480 kDa, and 1048 kDa, Waltham, MA, USA). Calibration was conducted by diluting 2.5 µL of the standard mixture directly into a 10 µL buffer drop placed on the illuminated glass coverslip. Isolated SMAD4 constructs and assembled complexes were prepared at 50–100 nM stock concentrations. A 1 µL aliquot of sample was homogenized into the buffer drop to yield final assay concentrations of 2.63 nM for SMAD4 constructs and 5.26 nM for SMAD3. On the OneMP system, isolated SMAD4 constructs (expected monomeric mass ∼38 kDa) yielded a Gaussian distribution centered at ∼50 kDa, reflecting the resolution limits of the instrument at low molecular weights.

To achieve higher resolution for low molecular weight species and nanobody-bound complexes, subsequent measurements were performed on the Refeyn TwoMP system. Mass calibration for TwoMP experiments was conducted using the MassFrence P1 protein standard (Refeyn Ltd.). The calibrant was diluted 1:250 in the assay buffer and mixed 1:1 with a pre-placed 10 µL buffer drop on the instrument (1:500 final dilution). High-resolution TwoMP measurements enabled clear discrimination between monomeric/dimeric species (27–65 kDa) and homotrimeric complexes (81–92 kDa). Complexes comprising SMAD3, SMAD4, and nanobody NbS4A7 were generated at a 2:1:1 stoichiometric ratio and measured at final concentrations of 10.5 nM SMAD3, 5.3 nM SMAD4, and 5.3 nM NbS4A7.

All measurements were acquired in standard acquisition mode using normal spatial scale settings. To prevent time-dependent dissociation or aggregation artifacts, each sample was prepared fresh immediately prior to drop dilution and measured under identical incubation timeframes across both OneMP and TwoMP runs. Data inclusion criteria were restricted to count rates between 80 and 150 events per frame, with ratiometric signal intensities exceeding 0.07. Mass histograms and fitted Gaussian distributions were generated and analyzed using DiscoverMP (Refeyn Ltd.) and PhotoMol software (<u>spc.embl-hamburg.de</u>).

### Surface plasmon resonance (SPR)

The dissociation kinetics of complexes formed between SMAD4 variants and R-SMADs were analyzed by surface plasmon resonance (SPR) using a Biacore T200 system (Cytiva). SMAD4 constructs were immobilized onto CM5 S Series sensor chips via standard amine coupling to target levels of 400–500 RU. Experiments were conducted at 25 °C at a flow rate of 50 µL/min (low sample consumption mode) in 20 mM Tris pH 7.5, 100 mM NaCl. Analyte (R-SMAD) concentrations ranging from 1.56 to 25 µM were injected over the surface with association and dissociation phases set to 120 s and 180 s, respectively. Surfaces were regenerated between cycles using 1.5 M NaCl, 0.5% Tween-20. To eliminate mass transfer effects arising from the complex monomer-dimer-trimer equilibria of R-SMAD species, only the dissociation phases were evaluated.

Data were processed using Biacore T200 Evaluation Software (Cytiva) and fitted to a 1:1 dissociation model using Equation 2, where R is the response signal (RU), R_0_ is the initial response at the onset of dissociation (t_0_ = 120 s), k_off_ is the dissociation rate constant (s^-1^), and Offset represents residual response above baseline post-dissociation. Dissociation rate constants were subsequently converted to complex half-lives (t_1/2_) using Equation 3.

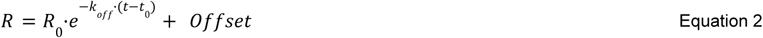

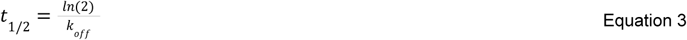

### Native gel electrophoresis

To assess protein-DNA interactions, electrophoretic mobility shift assays (EMSAs) were performed using Cy5-labeled DNA probes. Protein-DNA complexes were resolved on 1.5mm thick 4.5% polyacrylamide gels polymerized in Tris-Glycine buffer and visualized using a Typhoon imager (Cytiva). DNA duplexes were prepared by annealing complementary oligonucleotides. Protein-DNA binding reactions were carried out under optimized conditions, and samples were loaded onto native polyacrylamide gels. Gel electrophoresis was performed at 100V during 45 min. EMSA were performed at 4 °C to maintain protein-DNA native complex stability.

### ITC experiments

ITC measurements were performed using a nano ITC calorimeter (TA Instruments) at 37°C and 260 rpm. For SMAD complexes, 50 μL of 100 μM (variants) or 280 μM SMAD4 (WT) (314-552aa) (syringe) and 190 μL of 30 μM or 80 μM SMAD3MH2-DVD (cell) were used for variants and WT constructs, respectively. The titrant volume was divided into a total of 17 injections. Concentrations were re-measured after degassing the samples to account for possible changes due to evaporation using a NanoDrop system. All proteins were dialyzed overnight at 4 °C in the 20 mM Tris pH 7.5, 100 mM NaCl, 2 mM TCEP. Binding isotherms were analyzed using NanoAnalyze software (TA Instruments, New Castle, DE, USA) using the one site fitting. Baseline controls were performed with buffer and SMAD4 variants to subtract SMAD4 dilution effects on the measurement. Thermodynamic values are typically reported per mole in kJ/mol.

### X-ray crystallography

Crystallization conditions were screened at 4 and 20 °C, at the Automated Crystallography Platform (PAC) of IBMB-CSIC and IRB Barcelona (Barcelona, Spain) using a Phoenix Crystallization Robot (Art Robbins, Sunnyvale, CA, USA). The set up consisted of sitting-drop vapor diffusion experiments in the SWISSCI 96-well format 3-lens plates with 30 µL of the reservoir solutions. To minimize the impact of the crystallization conditions and temperature on our comparative analysis, particularly of the subunit-subunit interfaces, we selected for the comparison only structures that originated from very similar conditions, which were also alike for the WT protein we solved earlier (pdb_9hwe) <u>(Macias et al. 2025</u>. Initial phase determination was performed by molecular replacement with PHASER <u>(McCoy 2007)</u> using the WT structure as a template (pdb_9hwe). In the structures, the asymmetric unit (ASU) contains a single MH2 monomer that assembles into a canonical head-to-tail homotrimer within the crystal lattice. Because the residues involved in the inter-subunit contacts in the Smad4/Smad4 interface and in the SMAD4/R-SMAD interface are the same, we extrapolate the homotrimer contacts those of the heterotrimer context for variants lacking cryo-EM data, although certain parts of the helical bundle and, in particular, the C-terminal tail, could not be resolved in the crystals structures. The crystallization conditions were as follows:

- **I500V (314-552):** a 300 nL crystallization drop was generated by adding 100 nL of the purified protein at 7.5 g/L in the SEC buffer to 200 nL of a precipitant solution consisting of 25% PEG 3350, 0.2 M lithium sulfate, and 0.1 M Bis-Tris pH 6.5. Crystals grew at 20 °C and were cryo-protected by immersion in the reservoir solution supplemented with a 25% v/v cryo mixture made of 60% v/v glycerol and 40% v/v PEG 400.
- **I500T (314-552):** a 300 nL crystallization drop was generated by adding 150 nL of the purified protein at 10 g/L in the SEC buffer to 150 nL of a precipitant solution consisting of 20% PEG 3350 and 0.2 M lithium sulfate. Crystals grew at 20 °C and were cryo-protected by immersion in the reservoir solution supplemented with a 25% v/v cryo mixture made of 60% v/v glycerol and 40% v/v PEG 400.
- **I500L (314-552):** a 300 nL crystallization drop was generated by adding 150 nL of the purified protein at 8 g/L in the SEC buffer supplemented with 0.05 M lithium sulfate to 150 nL of a precipitant solution consisting of 25% PEG 3350, 0.2 M lithium sulfate, and 0.1 M HEPES pH 7.5. Crystals grew at 20 °C and were cryo-protected by immersion in the reservoir solution supplemented with 30% v/v glycerol.
- **I500M (314-552):** a 300 nL crystallization drop was generated by adding 150 nL of the purified protein at 10 g/L in the SEC buffer to 150 nL of a precipitant solution consisting of 20% PEG 3350 and 0.2 M sodium sulfate. Crystals grew at 20 °C and were cryo-protected by immersion in the reservoir solution supplemented with a 25% v/v cryo mixture made of 60% v/v glycerol and 40% v/v PEG 400.
- **R496C (314-552):** a 300 nL crystallization drop was generated by adding 150 nL of the purified protein at 8 g/L in the SEC buffer to 150 nL of a precipitant solution consisting of 25% PEG 3350, 0.2 M sodium sulfate, and 0.1 M HEPES pH 7.5. Crystals grew at 20 °C and were cryo-protected by immersion in the reservoir solution supplemented with a 25% v/v cryo mixture made of 60% v/v glycerol and 40% v/v PEG 400.
- **WT-P6_3_ (314-552):** a 200 nL crystallisation drop was generated by adding 100 nL of the purified protein at 10 g/L in the SEC buffer supplemented with 0.8 mM IP3 to 100 nL of a precipitant solution consisting of 20% PEG 3350 and 0.2 M sodium tartrate. Crystals grew at 20 °C and were cryo-protected by immersion in the reservoir solution supplemented with 20% v/v glycerol. No zinc or other metal salts were added before crystallization.

Metal-binding sites were modeled as Zn(II) based on coordination geometry, metal–ligand distances in the 2.06 Å crystal structure, and anomalous scattering analysis. Although data were not collected at a wavelength optimized for anomalous signal, anomalous differences at 0.979 Å produced strong anomalous Fourier peaks (13–20 σ) at both metal sites and enabled site-specific f″ refinement in Phenix <u>(Echols et al. 2014)</u>. Refinement yielded f″ ≈ 2.5 e⁻ per site, closely matching the theoretical value for Zn (2.48 e⁻) at 0.979 Å and exceeding those for Cu (2.20 e⁻) or Ni (1.95 e⁻). Upon cation binding, the two coordinating histidine nitrogens move from 3.6 Å apart in metal-free F4₁32 crystals to 2.9–3.3 Å in P36 crystals. The absence of this density in the wild-type cryo-EM reconstruction likely reflects higher DTT concentrations added prior to vitrification, as DTT chelates Zn(II) <u>(Cornell and Crivaro 1972)</u>.

The diffraction data were collected at the BL13-XALOC (ALBA synchrotron, Barcelona, Spain) and ID30A-3 (ESRF, Grenoble, France) beamlines. Data processing, scaling, and merging were performed with autoPROC <u>(Vonrhein et al. 2011)</u> with anisotropy correction by STARANISO <u>(Tickle et al. 2018)</u> at the ALBA synchrotron or the Crystallographic Information Management System (CRIMS) at the HTX lab, EMBL <u>(Cornaciu et al. 2021)</u>. The CC1/2 criterion was used for selecting the diffraction resolution cut-off <u>(Diederichs and Karplus 2013)</u>. Initial phase determination was performed by molecular replacement with PHASER <u>(McCoy 2007)</u>. Refinement and structure optimization were performed using phenix.refine <u>(Liebschner et al. 2019)</u> and PDB-REDO <u>(Joosten et al. 2014)</u> with REFMAC <u>(Murshudov et al. 1997)</u>. Manual model building was done in COOT <u>(Emsley et al. 2010)</u>. Hydrogen atoms were used in the refinement either in the normal mode (phenix.refine) or in the riding hydrogens mode (PDB-REDO). UCSF Chimera <u>(Pettersen et al. 2004)</u> and PyMol (The PyMOL Molecular Graphics System, Version 3.1.0 open source Schrödinger, LLC) were used for visualization, analysis, and figures.

For the re-refinement of the SMAD3/SMAD4 heterotrimer (PDB code 1U7F), the structure factors (with data up to 2.49 Å resolution) and the coordinates model (refined up to 2.60 Å resolution) were downloaded from the RSCB PDB (https://www.rcsb.org/structure/1U7F), the residues proximal to the mutation spots were remodeled in COOT to better fit (if applicable) the electron density maps, and such a remodeled coordinates file was refined using PDB-REDO against the downloaded coordinates file. The resulting pdb and mtz files are included as supplementary files. The re-refined Rwork/Rfree factors were 0.242/0.254, and the completeness was 91.9%.

### Interchain distances and Interface metrics

**Assembly.** Each crystal has one chain per asymmetric unit (two for WT P63; chain A was used). The trimer was generated with the crystallographic symmetry operators of each file (gemmi 0.7.5 (Wojdyr 2022) CCP4 suite) from the two symmetry mates whose Ser368 OG lies within 3.6 A of a Glu526 carboxylate oxygen of the reference subunit, and vice versa. Interchain distances (Supplementary Figure 4b). For each residue pair (Glu526-Ser368’, residue 496-Ser368’, Arg497-Glu377’, Asp493-Arg380’; primed residues belong to the neighbouring subunit) the distance is the shortest distance between side-chain polar heavy atoms (N, O, S) of the two residues, taking the closest symmetry mate in the crystal lattice. Backbone and hydrogen atoms were excluded; residue 496 is Arg (WT, I500 variants) or Cys (R496C), for Cys the SG atom was used. Interface metrics (Supplementary Table 2). Solvent-accessible surface was computed with the Shrake-Rupley algorithm (own implementation, 1.4 A probe, 960 points per atom, Bondi radii), on protein heavy atoms, using the first alternate conformation where present. Buried surface area (BSA) per subunit is the loss of solvent-accessible surface of one subunit on forming the interface with its neighbor. Because the interface has two non-equivalent faces, both values are reported. Contacts are heavy-atom pairs closer than 4 A across one interface. The subunit centre of mass (COM) separation is the mean of the three pairwise distances between subunit centroids of the trimer, and Rg is the radius of gyration of the trimer; both were calculated on the heavy atoms of the 173 residues modelled in all 13 crystals (320-415, 425-450 and 492-542), so that differences in modelled chain termini and loops do not contribute. Clustering. The 13 x 4 distance matrix (values in A, not scaled) was displayed as a heatmap. Rows (crystals) were clustered hierarchically with Pearson correlation distance (1 - r) and average linkage.

### Cryo-EM Sample preparation, Data Collection and Data Processing

Concentrated complex stock solutions were prepared at 15 μM NbS4A7, 15 μM SMAD4 MH2_272 (WT, I500V and R496C variants), and 45 μM SMAD3 MH2_DVD in 20 mM Tris pH 7.5, 200 mM NaCl, 1 mM DTT and incubated for at least one hour on ice prior to 4 μM dilution and plunge-freezing. For the WT, the solution was supplemented with dithiothreitol (DTT) to a 3 mM final concentration immediately before vitrification to prevent aggregation when contacting the HexAuFoil® grids (SPT Labtech) (WT complex) or QUANTIFOIL® gold grids (300 mesh, R 0.6/1.0 100 Holey Gold Supports, Grossloebichau, Germany) (variants). Grids were plasma-cleaned for 60 seconds with 90% argon and 10% oxygen in a NanoClean 1070 Plasma Cleaner (Fischione, Export, PA, USA). Vitrification was performed using a Vitrobot Mark IV (Thermo Fisher Scientific) with Whatman blotting paper (Cytiva, CAT No. 10311807) at 8 °C and 100% humidity, applying 3 μL of sample on each grid with a blot force of 5 and a blot time of 2 seconds. The grid was plasma-cleaned for 60 seconds with 90% Ar and 10% O_2_ in a NanoClean 1070 Plasma Cleaner (Fischione). Vitrification was performed using a Vitrobot Mark IV (Thermo Fisher Scientific) with Whatman grade 597 blotting paper (Cytiva, CAT No. 10311807) at 8 °C and 100% humidity, applying 3 μL of sample on the grid with a blot force of 5 and blot time of 5 seconds.

Data collection was performed at the EMBL Imaging Centre (Heidelberg, Germany) using SerialEM version 4.2.6 (Mastronarde 2003) on a Titan Krios G4 TEM operated at 300 kV equipped with an E-CFEG, a Selectris X energy filter, and a Falcon 4i direct electron detector (Thermo Scientific, Eindhoven, The Nederlands). For all three complexes, images were acquired at stage tilts of 0° and 30° to improve angular sampling, using a total electron fluence of 60 e⁻/Å².

The WT dataset was collected at a calibrated pixel size of 0.5726 Å, with movies recorded as 666 raw electron-event representation (EER) fractions. The I500V and R496C datasets were collected at calibrated pixel sizes of 0.934 Å and 0.731 Å, respectively. Dataset-specific movie counts, particle numbers, and processing workflows are shown in **Schematics 1 and 2**.

Movies were imported with an EER fractionation resulting in 60 frames/moves and an upsampling factor of 1 (4096×4096) into CryoSPARC v4.7.1 <u>(Punjani et al. 2017)</u>, to perform Patch Motion Correction, Patch CTF Estimation and Exposure curation. Particles were picked on the accepted micrographs with crYOLO 1.9.9 <u>(Wagner et al. 2019)</u> (WT, R496C) or the Cryosparc Blob Picker (I500V). These picks were cleaned with the Micrograph Junk Detector job. The remaining particles were extracted with an extraction box size of 416 pixels for the WT, 240 for I500V and 288 for R496C and Fourier cropped to 104, 80 and 72 pixels respectively. Successive rounds of 2D classification, 3D reconstruction, and 3D classification were used to select particles for high-resolution refinement. Selected particles were transferred to RELION v5.0 <u>(Scheres 2012)</u> with UCSF pyem <u>(Asarnow et al. 2019)</u> (https://github.com/asarnow/pyem). Movies were gain- and motion-corrected with dose weighting using MotionCor2 <u>(Zheng et al. 2017)</u>, (EER fractionation: 11, dose per frame: 1e/Å^2^, doing dose-weighting) and CTF-estimation with CTFFIND4 <u>(Rohou and Grigorieff 2015)</u>. For focused refinement, the best particles were sequentially 3D refined with blush regularization <u>(Kimanius et al. 2024)</u> after CTF-based refinement of defocus values, magnification anisotropy, and optical aberrations <u>(Zivanov et al. 2020)</u> and Bayesian polishing <u>(Zivanov et al. 2019)</u>, until no improvement was observed.

Initial atomic coordinates were generated with ModelAngelo with the initial NbS4A7 model being replaced with the NbS4A7 coordinates from the Smad4/NbS4A7 complex structure (pdb_00009qe8) <u>(Condeminas et al. 2026; Jamali et al. 2024)</u>. Refinement was performed with phenix.real_space_refine and manually optimized in Coot <u>(Liebschner et al. 2019; Emsley et al. 2010)</u>.

To characterize structural heterogeneity in the I500V complex, we performed 3D Variability Analysis (3DVA) in cryoSPARC using the particle subset indicated in the processing workflow (**Schematics 1**). The first three principal components of variability (eigenvectors; components 0–2) were analysed, and the corresponding map series were generated using 3D Variability Display. Videos were prepared in UCSF ChimeraX to examine variability in the SMAD4 C-terminal region and the N-terminal extensions of the SMAD3 MH2 domains (**Supplementary Video 1**).

## Data availability

The atomic coordinates and structural factors have been deposited in the PDB with the accession codes indicated in Tables 1 and 2. Data shown in the figures are included as Excel files in the Supplemental Information section.

## Acknowledgements

Special thanks go to the Myhre Syndrome patient associations for their support, to the staff at the EMBL image center for help, and to Christian O. Reyes, project director of Rec Bison Media SL, for helping bring visibility to this disease through a documentary work. We also thank the Protein Expression Core Facility (IRB Barcelona) for providing reagents, Marta Taulés from the CCiTUB for support with the SPR experiments, Dr. D. Derbyshire at the ProLinC facility (University of Linköping), Dr. Bernhard Brutscher at the IBS Grenoble, and the Automated Crystallography Platform (IBMB-CSIC), Drs. R Boer, F. Gil, X. Carpena, and I. Crespo at the ALBA and E. Mossou at the ESRF Synchrotrons, respectively, for assistance and support in using beamlines and to Dr. S. Mattei and his team at the Imaging Centre of EMBL Heidelberg for their support during M.C.’s internship.

## Funding

Research in M.J.M. lab is funded by the La Caixa health 2025 programme (HR25-00235) as well as the PID2021-122909NB-I00 grant from the Spanish Ministerio de Ciencia e Innovación MCIN/ AEI /10.13039/501100011033, and by the Next Generation EU/ PRTR and FEDER, A Way of making Europe. We also received support from the Agencia de Gestió d’Ajuts Universitaris i de Recerca, AGAUR (SGR 866-2021). Access to the ProLinC and SPC facilities was possible thanks to the Molecular-Scale Biophysics Research Infrastructure MOSBRI-2022-97 grant and by INSTRUCT funded by the EU Horizon 2020 (No 101004806). This work has been carried out as part of the canSERV project funded by the EU Horizon Europe programme (access to IBS NMR facility). We also acknowledge the funding provided to access ALBA (proposal 2021095308) and ESFR (proposal MX2356 ID30A-3 / MASSIF-3). We acknowledge the access and services provided by the Imaging Centre at the EMBL, supported by the Boehringer Ingelheim Foundation. R.J.K. was funded by the EU Horizon 2020 RI program, Marie Skłodowska-Curie COFUND actions and IRB Barcelona (IRB_DREAM no. 945352), and M.R.R. by a PhD AGAUR grant. CT and R.J.K. are currently founded by La Caixa Health 2025. Research in A.M. laboratory is funded by the Foundation K.G. Jebsen and The Neuro-SysMed Center (RCN, Project No. 288164), M.G.A. received funding from the European Union’s Horizon 2020 RI program under grant agreement No. 101004806.

We gratefully acknowledge institutional funding from the CERCA Programme of the Government of Catalonia, IRB Barcelona, the BBVA Foundation, the core facility BiSS and the department of Biomedicine, University of Bergen, EMBL (Grenoble and Hamburg), and the Spanish Ministry of Science and Education through the Centres of Excellence Severo Ochoa Awards. M.J.M. is an ICREA Programme Investigator.

This work is dedicated to all individuals living with Myhre syndrome and to their families.

